# Functional diversification despite structural congruence in the HipBST toxin-antitoxin system of *Legionella pneumophila*

**DOI:** 10.1101/2022.11.26.518041

**Authors:** Jordan D. Lin, Peter J. Stogios, Kento T. Abe, Avril Wang, John MacPherson, Tatiana Skarina, Anne-Claude Gingras, Alexei Savchenko, Alexander W. Ensminger

**Affiliations:** Department of Molecular Genetics, University of Toronto, Toronto, Ontario, M5G1M1, Canada; Department of Biochemistry, University of Toronto, Toronto, Ontario, M5G1M1, Canada; Lunenfeld-Tanenbaum Research Institute, Sinai Health, Toronto, Ontario, M5G 1X5, Canada; Department of Chemical Engineering and Applied Chemistry, University of Toronto, Toronto, M5S 3E5, Canada; Department of Microbiology, Immunology and Infectious Diseases, University of Calgary, Calgary, T2N 4N1, Canada; Center for Structural Genomics of Infectious Diseases (CSGID), University of Calgary, Calgary, T2N 4N1, Canada

**Keywords:** Legionella pneumophila, toxin-antitoxin system, HipBA, HipBST, Lpg2370, split-HipA, kinase toxin

## Abstract

Toxin-antitoxin (TA) systems are abundant genetic modules in bacterial chromosomes and on mobile elements. They are often patchily distributed and their physiological functions remain poorly understood. Here, we characterize a TA system in *Legionella pneumophila* that is highly conserved across *Legionella* species. This system is distantly related to *Escherichia coli* HipBST and we demonstrate that it is a functional tripartite TA system (denoted HipBST_Lp_). We identify HipBST_Lp_ homologs in diverse taxa, yet in the Gammaproteobacteria these are almost exclusively found in *Legionella* species. Notably, the toxin HipT_Lp_ was previously reported to be a pathogenic effector protein that is translocated by *L. pneumophila* into its eukaryotic hosts. Contrary to this, we find no signal of HipT_Lp_ translocation beyond untranslocated control levels and make several observations consistent with a canonical role as a bacterial toxin. We present structural and biochemical insights into the regulation and neutralization of HipBST_Lp_, and identify key variations between this system and HipBST_Ec_. Finally, we show that the target of HipT_Lp_ is likely not conserved with any characterized HipA or HipT toxin. This work serves as a unique comparison of a TA system across bacterial species and illustrates the molecular diversity that exists within a single TA family.

## INTRODUCTION

Bacterial genomes are a mosaic of genes that are highly conserved and often critical for growth (the core genome) or variable and typically non-essential (the accessory genome). The accessory genome is frequently comprised of genes acquired through horizontal exchange between cells, and the plasticity of this portion of the genome can promote organismal diversification while also maintaining robustness to changing selective pressures (1). The contribution of accessory genomic elements to bacterial physiology is therefore of critical importance to understand, in particular with regard to genes that are considered accessory but demonstrate patterns of significant conservation across species (2).

Toxin-antitoxin (TA) systems are abundant and widely distributed components of the accessory genome (3). They are typically bipartite genetic modules found in nearly all bacterial chromosomes—often in multiple types and copies—as well as on mobile DNA elements (4, 5). Despite their widespread occurrence, these genetic systems are typically poorly conserved across related strains and species (5, 6). Their function in bacterial physiology remains poorly understood, however current evidence supports the involvement of some systems in phage defense and DNA stabilization (7), while others appear to contribute to bacterial persistence (8). TA systems are broadly divided into eight major groups based on antitoxin activity and molecule type (7), with the most heavily studied being the type II systems. Type II TA systems are composed of a protein toxin that can reversibly arrest cellular growth and a protein antitoxin that neutralizes the toxin through direct physical interaction. The toxin is typically refractory to proteolytic degradation, whereas the antitoxin is often rapidly degraded by the cell and needs to be frequently replenished (9). This leads to cellular addiction to the TA system—a property which allows them to act as selfish elements and consequently, to stabilize mobile DNA (10).

Numerous mechanisms of cell toxification and toxin neutralization have been reported for type II TA systems (11). Genetic diversification has even been observed within the same TA system, such as the recent discovery of a split-kinase variant of the well-studied HipBA system in *Escherichia coli* (12). HipBA is a bipartite TA module composed of the HipA kinase toxin and HipB antitoxin, and has been implicated in regulating bacterial persistence (13). The newly described TA family HipBST is related to HipBA but displays a tripartite architecture: an N-terminal subdomain in the HipA kinase is encoded by a separate protein (HipS) and this functions as the antitoxin in the system (12). The catalytic core of HipA is expressed as a single protein (HipT) that retains toxin activity, while the canonical antitoxin HipB appears to transcriptionally regulate the locus. This unusual tripartite variant is therefore considerably divergent from the HipBA system, despite their relatedness, exemplifying both the undiscovered diversity within TA systems and their potential as substrates for molecular evolution.

Most knowledge of TA biology has been gleaned from studying a small number of bacterial species, thereby limiting the ability to characterize new and divergent modules. Furthermore, given their near ubiquity, modularity, and portability, it remains unclear whether homologous TA systems function similarly within different bacterial hosts (7). To address this, we sought to investigate the TA landscape within the intracellular pathogen *Legionella pneumophila*. *L. pneumophila* is a gram-negative environmental bacterium found within most global freshwater environments, where it parasitizes numerous protozoan species (14). To maintain its broad host range, *L. pneumophila* utilizes a large arsenal of translocated virulence proteins (termed ‘effectors’) to transform the host cell into a replicative niche (15). The genome of *L. pneumophila* contains the largest known assemblage of effectors (∼10% of genes), many of which contain homology to eukaryotic proteins and have likely been acquired via horizontal gene transfer from its hosts (16). *L. pneumophila* effectors are abundant in seven accessory genomic clusters of non-essential genes (17), and despite their critical role in pathogenesis, most are dispensable for growth in any one host (18). While the genome of *L. pneumophila* is enriched for effectors, it encodes only a small number of predicted TA systems (19) and is largely devoid of other mobile elements (such as prophage) which contribute to TA dissemination (7). We therefore wondered whether unique TA biology or functionality would be found in a species with the genomic constraints of an intracellular pathogen that has prioritized the acquisition of foreign genetic material from its hosts.

Here we report the characterization of a type II TA system in *L. pneumophila* that is highly conserved across *Legionella* species genomes. This *L. pneumophila* system is distantly related to the recently described HipBST module in *E. coli* (herein referred to as HipBST_Lp_ and HipBST_Ec_, respectively). Notably, the toxin HipT_Lp_ was previously reported to be a bacterial effector that is translocated into the eukaryotic host cell (20). Recently, a bi-functional role for HipT_Lp_ was also proposed, with activity in both the host and the bacterial cell (21). Contrary to this report, we demonstrate that HipT_Lp_ is not translocated at a level greater than what is observed for a negative (non-effector) control and has no obvious role within the eukaryotic cell. Instead, we show that HipBST_Lp_ encodes a functional tripartite TA system with clear effects on bacterial replication. Interestingly, HipBST_Lp_ represents a heretofore undiscovered subclade of HipBST systems that are widely distributed outside of the Gammaproteobacteria class but are almost exclusively found in the Legionellales order within that group. We demonstrate that the toxin HipT_Lp_ is a kinase and report a survey of its phosphoproteome within intoxicated cells. We find that despite their shared architecture, HipT_Lp_ does not appear to target the same cellular substrate as HipT_Ec_ or any characterized HipA toxin. We additionally present structural and biochemical insights into the regulation of HipBST_Lp_, including its mechanism of neutralization and points of divergence between HipBST_Lp_ and HipBST_Ec_. Overall, this work provides a unique comparison of a TA system across distant bacterial species and emphasizes that broad generalizations about system functionality should be made with caution. Instead, there appears to be considerable evolutionary diversity that remains underexplored within these abundant genetic elements.

## MATERIALS AND METHODS

### Strains and plasmids

All strains used in this study are listed in Table S1. All plasmids used in this study are listed in Table S2. All oligonucleotides used in this study are listed in Table S3. *L. pneumophila* strains used were derived from Lp01^JK (^22^)^, except for Lp02, which was used for the TEM-1 translocation experiments. *E. coli* XL1-Blue and TOP10 cells (Invitrogen) were used for cloning and plasmid maintenance, and BL21-GOLD (DE3) (Stratagene) cells were used for protein expression. TOP10 and BL21-GOLD (DE3) were additionally used for *in vivo* toxicity assays. *Saccharomyces cerevisia*e Y8800 (*MATa leu2-3,112 trp1-901 his3-200 ura3-52 gal4Δ gal80Δ GAL2-ADE2 LYS2::GAL1-HIS3 MET2::GAL7-lacZ cyh2R*) (23) was used for yeast two-hybrid assays and BY4742 (*MATα his3Δ1 leu2Δ0 met15Δ0 ura3Δ0*) (24) was used for *in vivo* toxicity assays. Bacterial expression plasmids were constructed using PCR products amplified from Lp01^JK^ genomic DNA with restriction cloning, or ligation-independent cloning for the pMCSG68-SBP-TEV protein purification constructs. PCR products were cloned into pDONR221-ccdB using BP clonase (Invitrogen), as per the manufacturer’s instructions. The resulting pDONR221 constructs were then used to clone the *hipBST_Lp_* genes into the yeast expression vectors pAG425GAL-ccdB (for yeast toxicity tests), pAG416GPD-ccdB, pDEST-DB-ccdB and pDEST-AD-ccdB (for the Y2H and Y3H assays) using LR clonase (Invitrogen) according to the manufacturer’s instructions. pAG425GAL-ccdB (Addgene plasmid # 14153; http://n2t.net/addgene:14153; RRID:Addgene_14153) and pAG416GPD-ccdB (Addgene plasmid # 14148; http://n2t.net/addgene:14148; RRID:Addgene_14148) (25) were gifts from Susan Lindquist. Vectors with the DNA-binding (DB) and transcription-activating (AD) domain of Gal4 (pDEST-DB, pDEST-AD) (26) were a kind gift from N. Yachie and F. Roth (University of Toronto, Canada). Genetic substitutions were introduced via site-directed mutagenesis using the Q5 site-directed mutagenesis kit (NEB) according to the manufacturer’s instructions. All generated constructs were confirmed by Sanger sequencing. Plasmids were introduced into *E. coli* via heat-shock transformation, into *S. cerevisia*e via the PEG/LiAc method (27), and into *L. pneumophila* via electroporation. The endogenous *hipBST* locus in *L. pneumophila* was deleted using the previously described MazF genome editing method (28) with minor modifications to produce scar-free, in frame deletions. Briefly, linear DNA containing a *mazF*-Kan^R^ identify those with kanamycin sensitivity. The resulting deletion mutant was screened by PCR using the oligos JL-P102 and JL-P103 (Table S3), and validated with Sanger sequencing using the oligo JL-P101.

### Media and culture conditions

Bacterial experiments and routine strain maintenance were performed at 37°C. *L. pneumophila* strains were grown in *N*-(2-acetamido)-2-aminoethanesulfonic acid (ACES)-buffered yeast extract and on charcoal AYE (CYE) agar plates supplemented with 0.4 g/L L-cysteine and 0.25 g/L ferric pyrophosphate. For liquid growth, cultures were inoculated from patches grown for 2 days. When required for selection or plasmid maintenance, media were supplemented with chloramphenicol (5 μg/mL), kanamycin (20 μg/mL), or thymidine (100 μg/mL). Ectopic gene expression was induced by isopropylthio-β-galactoside (IPTG; 100 μM) and repressed with glucose (1% v/v), unless otherwise indicated. *E. coli* strains were grown on lysogeny broth (LB, Miller) liquid media and agar. When required, media were supplemented with ampicillin (100 μg/mL), chloramphenicol (34 μg/mL), or kanamycin (40 μg/mL) for selection or plasmid maintenance. Ectopic gene expression was induced by arabinose (0.2% w/v) or IPTG (100 μM) and repressed with 1% glucose, unless otherwise indicated. Yeast experiments and routine strain maintenance were performed at 30°C. *S. cerevisia*e strains were grown on yeast peptone adenine dextrose (YPAD) medium (2% bacto peptone w/v, 1% yeast extract w/v, 2% glucose v/v, 180 mg/L adenine sulfate), or synthetic defined (SD) medium comprised of yeast nitrogen base with ammonium sulfate, supplemented with 2% glucose and all amino acids, lacking specific amino acids where necessary for selection or plasmid maintenance. When required, media lacking glucose were supplemented with galactose (2% v/v) to induce gene expression.

### TEM-1 β-lactamase translocation assays

Protein translocation was tested as described previously (29). Briefly, Lp02 strains carrying pXDC61 constructs containing the TEM-1 β-lactamase fused to a gene of interest were grown overnight to mid-log phase (OD_600_=1.5-2) and fusion protein expression was induced with IPTG (500 μM) for 3 hr. The cultures were then used to infect monolayers of differentiated U937 cells (RPMI 1640 medium, Gibco) in triplicate at various multiplicities of infection (MOI; 20, 50, 125) for 1 hr at 37°C with 5% CO_2_. LiveBLAzer dye (ThermoFisher K1095) was subsequently added to the cells and incubated for another 2 hr, followed by blue/green fluorescence detection using a TECAN Infinite M PLEX plate reader (blue fluorescence: 409 nm excitation, 460 nm emission, gain 140, integration time 20 μs; green fluorescence: 409 nm excitation, 530 nm emission, gain 140, integration time 20 μs). A blue/green fluorescence ratio > 1 was used as the threshold to determine translocation (30). Each experiment was performed in triplicate and repeated 3 times. Immunoblotting to confirm TEM-1 fusion protein expression was performed with anti-Beta Lactamase antibody (Abcam 12251).

### *In vivo* bacterial toxicity experiments

Growth assays were performed as follows: *E. coli* and *L. pneumophila* strains containing plasmids expressing the genes of interest were grown overnight in the presence of 1% glucose. Cultures were washed to remove glucose and adjusted to OD_600_ =0.1 in fresh media supplemented with either 1% glucose or arabinose/IPTG. Cultures were plated in triplicate in a flat bottom 96-well plate (100 μL volume), sealed with a Breathe-Easy sealing membrane (Diversified Biotech BEM-1), and optical density (600 nm) was monitored every 15 mins for 24 hr using an S&P growth curve robot. Each experiment was repeated a minimum of 3 times unless otherwise indicated.

### Protein expression and purification for *in vitro* experiments

Protein expression and purification were performed as described previously (31). Briefly, BL21-GOLD (DE3) cells carrying the desired His_6_-SBP-tagged constructs were grown at 37°C to OD_600_=0.5, at which time protein expression was induced with IPTG (1 mM) for 5 hr at 37°C. Cells were harvested by centrifugation (12,227 x *g* for 10 mins at 4°C), resuspended in lysis buffer (300 mM NaCl, 5% glycerol, 5 mM imidazole, 50 mM HEPES [4-(2-hydroxyethyl)-1-piperazineethanesulfonic acid] pH 7.5), lysed by sonication on ice (30% amplitude, 10 sec on, 10 sec off for 5 min) in the presence of 1 mM phenylmethylsulfonyl fluoride (PMSF), and the soluble fraction was obtained by high-speed centrifugation (34,957 x *g* for 20 mins at 4°C). The tagged proteins were purified by immobilized metal affinity chromatography using nickel-nitrilotriacetic acid beads. Columns were washed with wash buffer (300 mM NaCl, 30 mM imidazole, 15 mM HEPES pH 7.5, 5% glycerol), and eluted in 300 mM NaCl, 300 mM imidazole, 15 mM HEPES pH 7.5. Eluted protein was then dialyzed in 300 mM NaCl, 15 mM HEPES pH 7.5, 0.5 mM dithiothreitol (DTT), and concentrated using Vivaspin 5 kDa (HipB_Lp_, HipS_Lp_) and 10 kDa (HipT_Lp_) cutoff columns (GE Healthcare). Protein concentration was quantified using the Bradford assay according to the manufacturer’s instructions (BioShop). For additional purification (*in vitro* assays involving co-incubation), His_6_-SBP-tagged HipB_Lp_, HipS_Lp_, and HipT_Lp_ were injected into a Superdex S75 10/300 GL size exclusion column (GE Healthcare) equilibrated in dialysis buffer, and the purified fractions were collected.

### Crystallography and structure determination

The HipS_Lp_-HipT_Lp_ complex was crystallized as selenomethionine (Se-Met)-derivatized proteins and as native proteins (for the higher resolution final structure). Cloned in pETDuet vector, HipS_Lp_ and His_6_-HipT_Lp_ proteins were expressed in *E. coli* BL21(DE3)-Magic cells. For the (Se-Met)-derivatized protein complex, cells were grown in M9 minimal media (Shanghai Medicilon) with 1mM IPTG induction at 20°C when the OD_600_ reached 1.2. The native protein complex was expressed in ZYP-5052 auto-inducing complex medium (32) by incubating for a few hr at 37°C followed by overnight growth at 20°C. Overnight cell culture was then collected by centrifugation at 6,000 × *g* for 25 min at 4°C. Cells were resuspended in a binding buffer (100 mM HEPES [pH 7.5], 500 mM NaCl, 5 mM imidazole, and 5% glycerol [vol/vol]).The purification was performed as described above, which resulted in formation of the HipS_Lp_-HipT_Lp_ complex as judged by SDS-PAGE. HipB_Lp_ was expressed as native protein and purified as described above. Crystals were grown at room temperature (RT) using the vapor diffusion sitting drop method. For the HipS_Lp_-HipT_Lp_ (Se-Met) complex, 17 mg/mL protein was mixed with reservoir solution (0.1 M potassium chloride, 10 mM Tris pH 7, 20% (v/v) PEG 4K). For the HipS_Lp_-HipT_Lp_ (native) complex, 8 mg/mL protein was mixed with reservoir solution (0.1 M HEPES pH 7.5 and 30% (v/v) PEG 1K). For the HipB_Lp_ crystal, 20 mg/mL protein was mixed with reservoir solution (0.2 M ammonium sulfate, 0.1 M sodium acetate pH 4.6 and 25% (v/v) PEG4K). Crystals were cryoprotected with paratone oil. For the HipS_Lp_-HipT_Lp_ complex crystal, diffraction data at 100 K were collected at beamline 19-ID of the Structural Biology Center at the Advanced Photon Source, Argonne National Laboratory. For the HipB_Lp_ crystal, diffraction data at 100 K were collected at a home source Rigaku HF-007 rotating anode with Rigaku R-AXIS IV image plate detector. All diffraction data were processed using HKL3000 (33). The HipS_Lp_-HipT_Lp_ (Se-Met) complex was solved using Phenix.Autosol (34) which identified 7 Se-Met sites out of 7 total methionine residues in the asymmetric unit; the native complex was solved by Molecular Replacement using the Se-Met complex structure. The structure of HipB_Lp_ was solved by Molecular Replacement using the CCP4 online Balbes server and the structure of HipB_Ec_ (PDB 2WIU). All model building and refinement was performed using Phenix.refine and Coot (35). Atomic coordinates have been deposited in the Protein Data Bank with accession codes 8EZR, 8EZS and 8EZT.

### *In vitro* kinase assays

Purified recombinant His_6_-SBP-tagged HipB_Lp_, HipS_Lp_, and HipT_Lp_ (WT, D197Q, D219Q) were tested for ATP hydrolytic activity using the ADP-Glo kinase assay (Promega) per the manufacturer’s instructions. Briefly 1 µg of each protein was assayed in 25 µL reactions containing 40 mM Tris-HCl pH 7.5, 20 mM MgCl_2_, 0.1 mg/mL bovine serum albumin, 1 mM DTT, and 25 µM ATP. When included, the kinase inhibitor 5’-fluorosulfonylbenzoyl-5’-adenosine (FSBA) was supplemented to a final concentration of 20 mM. Reactions were incubated at 37°C for 30 min and luminescence was measured using a TECAN Infinite M PLEX plate reader (500 ms integration time). Each experiment was replicated independently a minimum of 2 times.

### Electrophoretic mobility shift assays

Purified recombinant His_6_-SBP-tagged HipB_Lp_, HipS_Lp_, and HipT_Lp_ were incubated with DNA comprising the promoter region (upstream 200 bp amplified with oligos JL-P355 and JL-P342) of the *hipBST*_Lp_ operon in binding buffer (400 mM KCl, 150 mM HEPES pH 7.5, 10 mM EDTA, 50% glycerol, 5 mM DTT) and incubated for 30 min at RT. The protein-DNA mixtures were then run on a 6% polyacrylamide gel with 1x TAE (Tris-acetate-EDTA) running buffer, stained with SYBR Green (Invitrogen), and imaged. Each experiment was repeated a minimum of 3 times.

### Yeast two-hybrid assays

Yeast two-hybrid (Y2H) experiments were performed as described previously (36). Briefly, proteins of interest were fused to either the GAL4 transcriptional activating domain (AD) or DNA-binding domain (DB), using the pDEST-AD-ccdB and pDEST-DB-ccdB constitutively active Gateway destination plasmids. Three independent clones of Y8800 containing pDEST-AD and pDEST-DB encoding gene fusions of interest were grown overnight at 30°C in liquid SD -Leu/Trp media supplemented with 2% glucose. These cultures were then stamped on plates containing (control) or lacking histidine (physical interaction selection). When required, a third protein was expressed constitutively from the pAG416GPD plasmid. Y8800 strains containing pDEST-AD, pDEST-DB and pAG416GPD together were grown with SD-Leu/Trp/Ura media supplemented with 2% glucose. Plates were imaged after 2 days of growth at 30°C. Each experiment was repeated a minimum of 2 times.

### Yeast spotting assays

Yeast spotting experiments were performed as described previously (36). Briefly, three independent clones of BY4742 containing the gene of interest in the pAG425GAL expression vector were grown in triplicate overnight at 30°C in liquid SD -Leu media supplemented with 2% glucose and adjusted to OD_600_=1 the following day. Five-fold serial dilutions of each culture were then prepared in a 96-well plate at a volume of 120 µL and stamped onto solid SD -Leu media supplemented with either glucose (2%; suppression) or galactose (2%; expression) using a VP 407AH pin tool (VP Scientific). Plates were imaged after 2 days of growth at 30°C. Each experiment was repeated a minimum of 3 times.

### ASKA genomic library rescue screen

The *E. coli* ASKA library (37) was prepared from glycerol stocked strains, pooled to a final concentration of 100 ng/μL, and 150 ng of the pool was electroporated into BL21-GOLD (DE3) cells containing *hipT_Lp_* cloned into the pCDF1-b expression vector. Cells were recovered for 1 hr at 37°C and transformations were plated on solid media containing either IPTG (100 μM) for gene expression or 1% glucose for repression and quantifying transformation efficiency. As a control, the empty vector pCA24N was transformed in an equivalent manner to the pooled library. Library transformations without gene expression yielded between 10^4^ and 10^5^ cells and library transformations were performed a minimum of three times for each experiment. Individual screening experiments were repeated 3 times. Transformants that grew under selective conditions were re-struck on selective media to ensure a stable rescue phenotype. Plasmids from the resulting strains were then prepared and Sanger sequenced using the oligo JL-P206.

### HipT_Lp_ *in vivo* expression for phosphoproteomic analysis

Cultures of *L. pneumophila* Δ*hipBST* carrying either pJB1806 or pJB1806::*hipT_Lp_* were inoculated from fresh patches and grown at 37°C in the presence of 0.5% glucose until mid-log phase (OD_600_ = 1.5-2). Expression was then induced with IPTG (100 μM) and 75 mL of cells were harvested prior to and 75 min post induction by centrifugation and flash freezing. Lysates were prepared by resuspending cell pellets in 2 mL NP-40 buffer (50 mM Tris-HCl pH 7.8, 150 mM NaCl, 1% NP-40, 1 mM PMSF, 1x PhosSTOP [Roche]), sonication on ice (5s on/10s off for 2 mins at 40% amplitude), and high speed centrifugation (13,000 x *g* for 30 mins at 4°C). Each experiment was repeated 2 times.

### In-solution protein digestion

Cell lysates from the Δ*hipBST* cultures (vector induced, HipT_Lp_ uninduced, and HipT_Lp_ induced) were quantified for total protein (Pierce BCA Protein Assay Kit, Thermo Scientific), and 2.5 mg of total protein from each sample was used. Samples were reduced with 5 mM DTT (37°C, 1 hr) and alkylated with 10 mM iodoacetamide (25°C, 45 min). Samples were digested for 2 hr at 25°C with constant agitation with a Trypsin/LysC mix (V5071, Promega), and overnight at 25°C with proteomics-grade porcine trypsin (T6567, Sigma-Aldrich). For both digestions, 6.25 µg of trypsin was used. Samples were desalted with 50 mg capacity Sep-Pak C18 cartridges (Waters Corporation). In brief, samples were acidified with trifluoroacetic acid (TFA) to a final concentration of 1% and centrifuged at 21,000 x *g* to remove precipitate. Columns were conditioned by passing one column volume (CV) of 100% acetonitrile (ACN), one CV of 50% ACN/0.1% formic acid (FA) and 4 CV of 0.1% TFA. Samples were loaded and desalted with 1% TFA, 1% FA, and washed peptides were eluted with 50% ACN/0.1% FA and concentrated by vacuum centrifugation.

### Dimethyl labelling

Samples were labelled as described previously (38) with some modifications. Peptides were resuspended in 50 mM HEPES pH 8.0, and the HipT_Lp_ uninduced, vector induced, and HipT_Lp_ induced samples were designated to be labelled with the light, intermediate and heavy channels, respectively. Peptides were mixed with freshly prepared CH_2_O (light, +28.0313 Da), CD_2_O (intermediate, +32.0564 Da) and ^13^CD_2_O (heavy, +36.0757 Da), and all solutions were made to 4% v/v. NaBH_3_CN (0.6 M) was added to the light and intermediate channels, and NaBD_3_CN (0.6 M) was added to the heavy channel. Samples were labelled at RT for 2 hr, quenched with FA (5% final) and desalted with Sep-Pak C18 Cartridges. Labelled peptides were quantified (Pierce Colorimetric Peptide Assay Kit, Thermo Scientific) and mixed 1:1:1 based on peptide amount.

### Phosphopeptide enrichment

Labelled, mixed peptides were phosphoenriched with TiO_2_ magnetic beads according to manufacturer’s protocols (MagReSyn). In brief, beads (100 µl) were equilibrated by three washes with loading buffer (0.1 M lactic acid in 80% ACN/5% TFA) before beads were incubated with 480 µg of peptides for 20 min at RT with end-over-end mixing. Samples were washed with 100 µl loading buffer, 80% ACN/1% TFA, and 10% ACN/0.2% TFA. Phosphopeptides were eluted twice in 1% NH_4_OH and acidified with FA to a final concentration of 2.5%. Samples were vacuum centrifuged and desalted with C18 stagetips. Stagetips were activated with 100% ACN/0.1% FA (buffer B) and washed twice with 5% FA. Samples resuspended in 5% FA were passed through each tip and washed with 0.1% FA (buffer A). Samples were eluted with a 2:1 mixture of buffer B and buffer A, and eluate was vacuum centrifuged until dry.

### LC-MS/MS

For data-dependent acquisition (DDA) LC-MS/MS, labelled, phoshoenriched peptides were analyzed using a nano-HPLC coupled to MS. Coated nano-spray emitters were generated from fused silica capillary tubing (75 µm ID, 365 µm OD) with a 5-8 µm tip opening, using a laser puller (Sutter Instrument Co., model P-2000). Nano-spray emitters were packed with C18 reversed-phase material (Reprosil-Pur 120 C18-AQ, 3 µm) resuspended in methanol using a pressure injection cell. The sample in 5% FA was directly loaded at 400 nl/min for 20 min onto a 75 µm x 15 cm nano-spray emitter. Peptides were eluted from the column with an ACN gradient generated by an Eksigent ekspert™ nanoLC 425, and analyzed on an Orbitrap Fusion Lumos Tribrid mass spectrometer (ThermoFisher). The gradient was delivered at 200 nl/min from 2.5% ACN with 0.1% FA to 35% ACN with 0.1% FA using a linear gradient of 120 min. This was followed by an 8 min gradient from 35% ACN with 0.1% FA to 80% ACN with 0.1% FA. After, there was an 8 min wash with 80% ACN with 0.1% FA, and equilibration for another 23 min to 2.5% ACN with 0.1% FA. The total DDA protocol was 180 min. The MS1 scan had an accumulation time of 50 ms within a mass range of 400–1500 Da, using orbitrap resolution of 120000, 60% RF lens, AGC target of 125% and 2400 volts. This was followed by MS/MS scans with a total cycle time of 3 seconds. Accumulation time of 50 ms and 33% HCD collision energy was used for each MS/MS scan. Each candidate ion was required to have a charge state from 2-7 and an AGC target of 400%, isolated using orbitrap resolution of 15,000. Previously analyzed candidate ions were dynamically excluded for 9 seconds.

### MS data processing and analysis

Data files were processed and searched using MaxQuant (version 2.1.4.0) querying against *Legionella pneumophila* sequences (RefSeq accession: GCF_001941585.1) that were modified to include common contaminants and reverse sequences (FragPipe v.18.0). Search parameters were set to search for tryptic cleavages allowing two missed cleavages. Variable modifications included methionine oxidation; protein N-terminal acetylation; asparagine deamidation; serine, threonine & tyrosine phosphorylation and lysine dimethylation. Fixed modifications included carbamidomethylation. A maximum of 5 modifications per peptide was allowed. Other settings were left as default parameters. For dimethyl labelling, light labels were processed with DimethLys0/Nter0, intermediate labels with DimethLys4/Nter4 and heavy labels with DimethLys8/Nter8. Biological duplicates were processed separately by assigning different Parameter Groups with PTM set to TRUE. Peptides that were flagged as contaminants, not labelled as phosphorylated or only found in one biological replicate were filtered out, and the intensity columns (e.g. Intensity.H) were used for analysis. Data analysis and plotting were conducted in R (version 4.2.0).

### HipBST conservation and non-essential cluster association search across *Legionella* species

Genome assemblies for 58 representative *Legionella* species were retrieved in July, 2020 from the NCBI RefSeq database (Table S4). When possible, alternative genome assemblies with higher sequencing coverage (39) were used instead of the representative genome. For *Legionella pneumophila*, a recently reannotated genome (Refseq ID: GCF_001941585.1) was used. Open reading frames (ORFs) were predicted using Prodigal (40) and homologs were determined using OrthoMCL (41). A core genome phylogeny was constructed by aligning all ORFs conserved across all species with MUSCLE and building a phylogenetic tree with RAxML (JTTDCMUT model selected by PROTGAMMAAUTO) (42). The resulting tree was annotated to show HipBST system presence across *Legionella* species using the R package ggtree (43). All software programs were run with default parameters. To identify non-essential gene clusters across *Legionella* species, genes in *L. pneumophila* were first annotated as essential if they were reported to demonstrate a growth defect in broth (17, 18) or replication within a host (18) when deleted, or were conserved across all 58 species. These annotations were then applied to homologous groups across all *Legionella* genomes. To predict non-essential clusters, a 10-gene sliding window analysis was performed on each genome and used to define regions where the local maximum of essential genes never exceeded 10%. In addition, each region was required to be larger than 1% of the total genome size.

### Phylogenetic analysis of HipBST systems across bacterial species

The NCBI RefSeq database was queried using cblaster (default parameters) (44) in November, 2021 to retrieve homologous sequences of HipBA or HipBST systems. We excluded systems that have an intergenic distance of above 150 bp and/or contained partial protein sequences. Five seed templates were used for this search: *E. coli* K12 HipBA (HipBA_Ec_; NP_416025.1, NP_416024.1), *Shewanella oneidensis* MR-1 HipBA (HipBA_So_; AAN53783.2, AAN53784.1), *Bacteroides uniformis* HipBA (HipBA_Bu_; WP_149924066.1, WP_149924064.1), *E. coli* O127 HipBST (HipBST_Ec_; WP_000563102.1, WP_001346664.1, WP_001262465.1), and *L. pneumophila* HipBST (HipBST_Lp_; AAU28429, AAU28430, AAU28431). The resulting protein sequences were clustered using MMseqs2 (45) to identify sequence-level representatives (80% identity, 80% coverage, coverage mode=1), aligned with MAFFT (E-INS-i) (46), and trimmed with trimAl (automated1 for parameter selection) (47). HipB sequences were aligned separately and the HipS/HipT sequences were concatenated prior to alignment with the HipA sequences. A HipBA/HipBST phylogeny was inferred via the maximum likelihood method using IQ-TREE (48). We used the edge-proportional partition model and performed partitioned phylogenetic analysis by segregating the protein sequence alignments into three regions: the HipB alignments, the HipA N-terminus/HipS alignments, and the HipA C-terminus/HipT alignments. Branch confidence was assessed using ultrafast bootstrap with 1000 replications. To explore the phylogenetic distribution of bacteria harbouring each system, species-level representatives of every taxon containing a system homolog from the initial search results were used to retrieve a phylogenetic tree from TimeTree (49). A second tree was retrieved after filtering for species with complete genome assemblies according to the NCBI Microbial Genome database, to ensure the absence of HipBA/HipBST systems is not due to genome incompleteness. Presence or absence were then compared for each system type across species and annotated on the resulting phylogenies using the R package ggtree (43).

## RESULTS

### *Legionella pneumophila* encodes a highly conserved HipBST locus

The genome of *L. pneumophila* has seven predicted TA loci in the Toxin-Antitoxin Database (19)—a comparatively small number relative to most other studied bacterial species. We first investigated the distribution of these systems across 58 representative species of *Legionella* in the NCBI Refseq database (Table S4) to determine whether any showed a signature of high conservation. Surprisingly, despite the mobile nature of TA systems and their patchy distribution within even closely related taxa (7), we identified one predicted system (*lpg2368-70*) that was found in nearly half of the *Legionella* species genomes (28/58 species; Figure 1A). In addition to these 28 species, eight others had homologs of individual genes from this locus (Figure 1A). This level of conservation is quite unusual for TA systems—particularly across different species genomes—and was striking given the considerable genomic diversity within this genus (50). Even effectors, which constitute a large portion of the *Legionella* accessory genome and are critical to pathogenicity, are poorly conserved across *Legionella* species (39, 50). Interestingly, *lpg2368-70* is similar to the well-characterized HipBA module (51), but possesses an atypical tripartite architecture in which the HipA homolog is split into two separate open reading frames (Figure 1B). Given its substantial conservation across *Legionella* species (>80% sequence similarity across homologs) and unusual genetic organization, we chose to investigate this putative TA system and its functionality in *L. pneumophila*.

**Figure 1.**
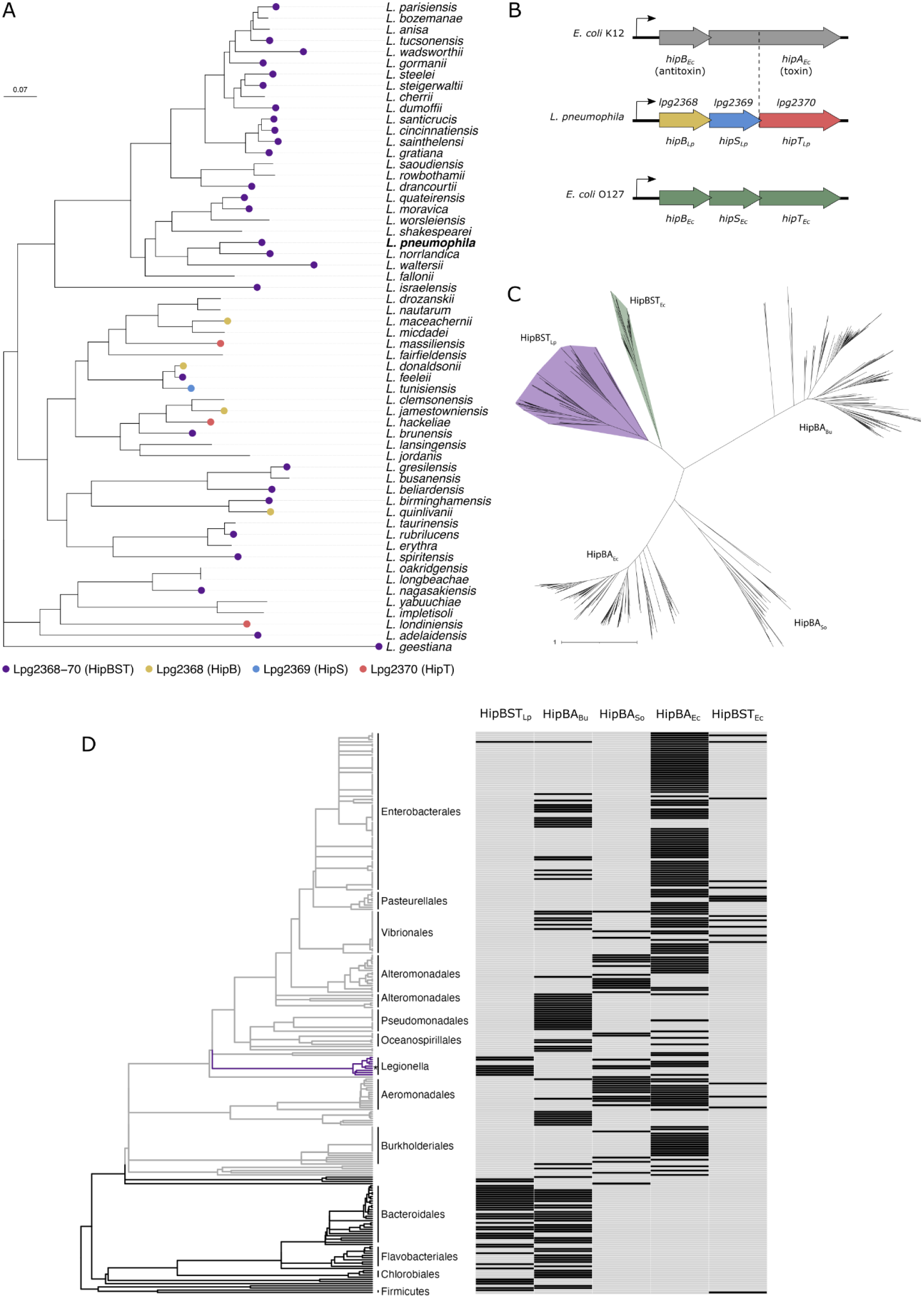
*Legionella* species contain a highly conserved and taxonomically restricted HipBST TA system. **(A)** A core genome phylogeny of 58 representative species in the *Legionella* genus (Table S4) showing conservation of Lpg2368-70 (HipBST_Lp_) homologs. The *Legionella* core genome was determined using OrthoMCL (41) to identify ORFs with conservation across all 58 species. Amino acid sequences were aligned with MUSCLE (41, 74), a phylogenetic tree was constructed with RAxML (42), and the tree was rooted with *Legionella geestiana* as the outgroup based on previous phylogenies (39, 50). Tree annotation and visualization were performed with ggtree (43). The scale bar denotes substitutions per site. **(B)** Schematic of the HipBA_Ec_, HipBST_Lp_, and HipBST_Ec_ TA systems. Lpg2368 has homology to the canonical antitoxin HipB, whereas Lpg2369 and Lpg2370 align to the N and C termini of the HipA toxin, respectively. **(C)** An unrooted phylogeny of HipBA and HipBST homologs retrieved from the NCBI Refseq database. Homology searching was performed using cblaster (44) with seed sequences from five different TA systems: *E. coli* K12 HipBA (HipBA_Ec_; NP_416025.1, NP_416024.1), *Shewanella oneidensis* MR-1 HipBA (HipBA_So_; AAN53783.2, AAN53784.1), *Bacteroides uniformis* HipBA (HipBA_Bu_; WP_149924066.1, WP_149924064.1), *E. coli* O127 HipBST (HipBST_Ec_; WP_000563102.1, WP_001346664.1, WP_001262465.1), and *L. pneumophila* HipBST (HipBST_Lp_; AAU28429, AAU28430, AAU28431). Retrieved sequences were clustered using MMseqs (45) to identify sequence-level representatives, aligned with MAFFT (46), and trimmed with trimAI (47). The phylogeny was constructed using IQ-TREE (48) and visualized with iTOL (75). The scale bar denotes substitutions per site. Raw and sequence-level representative hits are provided in Table S5. **(D)** Distribution of HipBA and HipBST TA systems across diverse bacterial taxa. A bacterial phylogeny was retrieved from TimeTree (49) for species containing at least one system in our homology search (Table S6). These were filtered to remove species without a complete genome, to ensure that genome incompleteness did not influence system detection. The presence of a homolog for each system is indicated for each species and systems are ordered by similarity of taxonomic distribution. The Pseudomonadota phylum is coloured light grey, the *Legionella* genus is coloured purple, and *L. pneumophila* is indicated with an asterisk.

The *lpg2368-70* locus encodes three genes within a predicted operon (Figure 1B) (52), with each ORF overlapping by 4 bp. The latter two genes were originally annotated as having homology to domains of the HipA toxin; the proteins encoded by *lpg2369* and *lpg2370* align with the N-terminus and C-terminus of HipA respectively. The third protein encoded by *lpg2368* has homology to the antitoxin HipB. These observations led us to hypothesize that Lpg2368-70 constituted a split-kinase TA system that was distantly related to HipBA. During the course of our studies, this was confirmed by the discovery of a similar system in *E. coli* (denoted HipBST_Ec_) (12). From this point forward, we refer to Lpg2368-70 as HipBST_Lp_. Despite its similar architecture, HipT_Lp_ bears only remote sequence similarity to either HipA_Ec_ (∼28% identity) or the newly characterized HipT_Ec_ (∼29% identity) (Figure S1A). This may explain why the HipBST_Lp_ homologs in *Legionella* species were not detected in previous searches for HipBA and HipBST systems (51). While they exhibit broad differences at the sequence level, the core catalytic residues in HipA_Ec_ and HipT_Ec_ are conserved in HipT_Lp_ (Figure S1A). The HipBST system of *L. pneumophila* therefore provided the opportunity to compare two variants of the newly described HipBST family.

### The HipBST_Lp_ system is phylogenetically and taxonomically distinct from HipBST_Ec_

As HipBST_Lp_ was not found in previous searches for HipBA/HipBST homologs, we sought to determine whether related modules were also undiscovered in other bacterial genomes and how these distant homologs fit into the broader Hip phylogeny. We queried the NCBI Refseq database for HipBA and HipBST homologs using five diverse systems as seed sequences: HipBA from *E.coli* K-12 (HipBA_Ec_), HipBA from *Shewanella oneidensis* (HipBA_So_), HipBA from *Bacteroides uniformis* (HipBA_Bu_), HipBST from *E. coli* O127 (HipBST_Ec_), and HipBST from *L. pneumophila* (HipBST_Lp_). While HipBA_Ec_ and HipBA_So_ have been characterized previously (51, 53), HipBA_Bu_ was included as a seed in order to further expand our search space, as this locus was found in close proximity to a HipBST_Lp_ homolog and was itself quite divergent from the other HipA/HipT homologs (Figure S1B). From this search, we identified 164 HipBA_Ec_, 37 HipBA_So_, 184 HipBA_Bu_, 31 HipBST_Ec_, and 86 HipBST_Lp_ sequence-level representative systems (Table S5), and used these sequences to construct a HipBST/HipBA phylogeny. Despite their shared tripartite architecture, the HipBST_Lp_ homologs cluster in a separate subclade from the HipBST_Ec_ sequences (Figure 1C), indicating system-level divergence even within the HipBST TA family. We discovered numerous HipBST_Lp_ homologs that are only distantly related to the 48 previously reported sequences (51), thereby expanding the sequence space of HipBST systems. In fact, the majority (60/86) of our HipT_Lp_ sequence-level representative sequences share less than 60% identity with any of the 48 sequences previously identified. The homologs of HipBA_Ec_ and HipBA_So_ also cluster separately, possibly as a consequence of sequence divergence underlying their different tertiary structures (53, 54), while the homologs of HipBA_Bu_ themselves form a clade distinct from all other systems (Figure 1C).

We next examined whether the various HipBA/HipBST homologs displayed any pattern of taxonomic distribution. To this end, we mapped the presence or absence of each system type onto a phylogeny of species that contained at least one system from our search and had a complete genome assembly in the Refseq database (Table S6). From this, we observed that HipBST_Lp_ homologs in the Gammaproteobacteria class are almost exclusively found within the Legionellales order, whereas most detected tripartite systems in this class are instead homologs of HipBST_Ec_ (Figure 1D, S1C). In contrast to their restricted distribution within the Gammaproteobacteria, HipBST_Lp_ homologs are widely distributed within distant taxonomic groups, particularly the FCB (Fibrobacterota, Chlorobiota, and Bacteroidota) superphylum. Interestingly, while HipBA_Bu_ homologs are prevalent throughout the bacterial phylogeny, homologs of HipBA_Ec_ and HipBA_So_ appear to be mostly restricted to the Pseudomonadota phylum. In summary, we used the previously undetected HipBST_Lp_ system to broaden the sequence space of known HipBST homologs. We observed that HipBST_Lp_ and HipBST_Ec_ sequences constitute distinct HipBST subclades, with HipBST_Lp_ homologs predominantly found in the Legionellales order within the Gammaproteobacteria class. This search therefore reveals considerably more genetic and taxonomic diversity within the HipBST TA family.

### HipT_Lp_ is not a translocated *L. pneumophila* effector

It was previously reported that HipT_Lp_ is translocated by *L. pneumophila* into its eukaryotic host via its Type IV secretion system (T4SS) (20), raising the intriguing possibility that it may represent a bacterial effector with an as-of-yet undefined host target. Notably, this hypothesis rests upon the report of a barely detectable signal in the TEM-1 translocation assay (21), in which TEM-1 β-lactamase is fused to a protein of interest and expressed in *L. pneumophila* cells infecting monolayers of differentiated U937 cells (29). Translocation is determined by monitoring emission from a fluorescent substrate within the host cells, which shifts from emitting green fluorescence to blue fluorescence upon cleavage of an internal beta-lactam ring by the TEM-1 fusion. Given the low level of translocation previously reported for HipT_Lp_ ^(^21^)^, we sought to confirm this observation. Using identical methodology, we observed that the translocation signal of HipT_Lp_ is equivalent to that of the negative control FabI—a cytosolic protein with no function in the host (Figure 2A). We observed the same negative results at several different multiplicities of infection (MOIs) and confirmed fusion protein expression by immunoblotting (Figure S2). To test whether the toxicity of HipT_Lp_ was impacting its observed level of translocation, we additionally tested two substitutions in predicted catalytic residues (D197Q, D219Q) that are conserved with HipA_Ec_ (Figure S1A) (55). However, no translocation signal above control levels was detected for these mutants (Figure 2A) despite their increased expression levels (Figure S2). These findings, along with the low level of translocation reported previously (21), do not support a role for HipT_Lp_ within the eukaryotic host. Further arguing against its role as an effector, we observe that overexpression of HipT_Lp_ does not display any growth inhibitory effects in yeast (Figure 2B), which is a common phenotype of effector expression (56). This is consistent with our identification of HipBST_Lp_ homologs within non-pathogenic bacterial species (Figure 1D, S1C) and argues for a canonical role with effects that are restricted to the bacterial cell.

**Figure 2.**
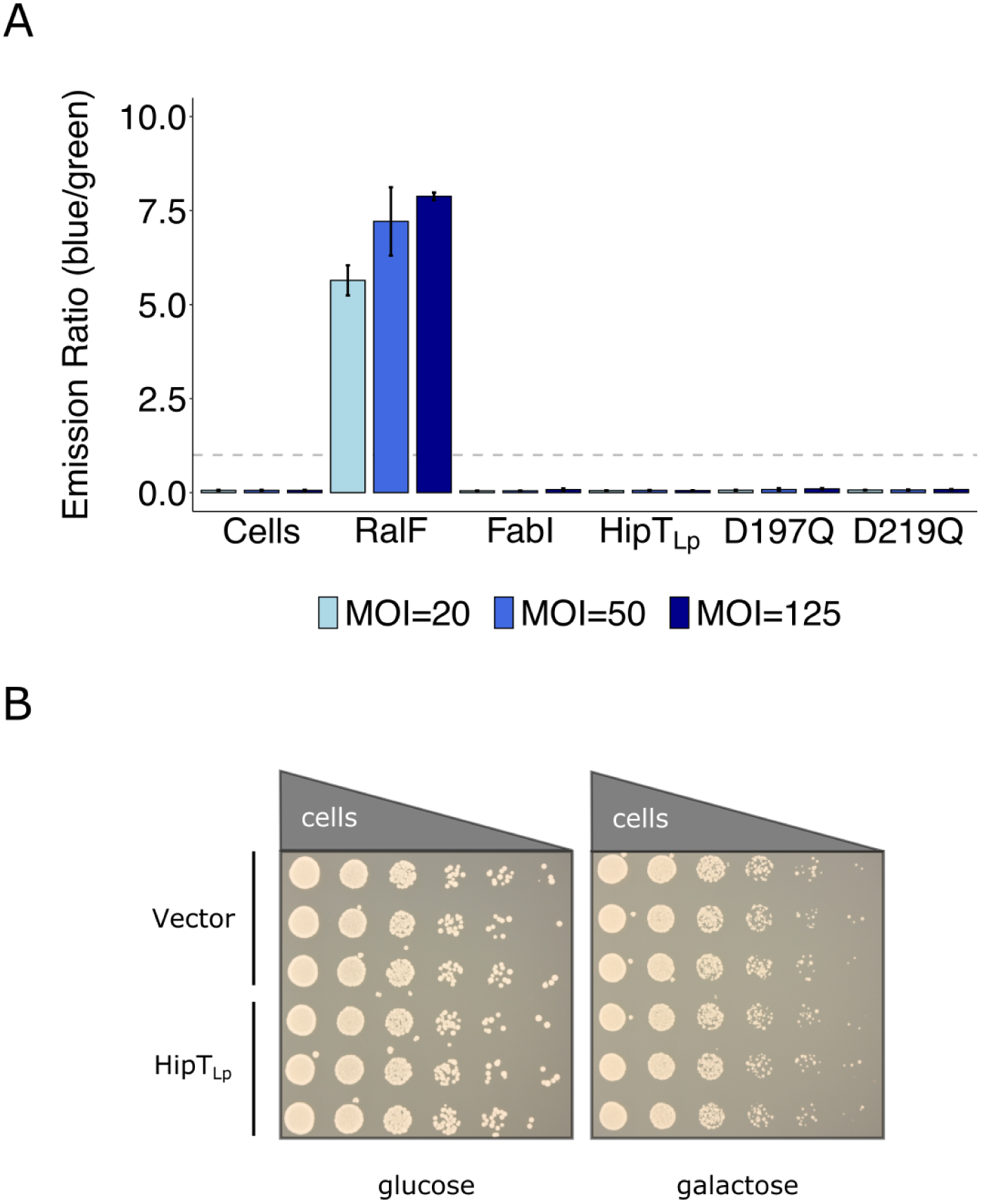
HipT_Lp_ is not a translocated bacterial effector. **(A)** TEM-1 β-lactamase translocation assays were performed with Lp02 cells infecting monolayers of differentiated U937 cells. Uninfected cells were used to determine background fluorescence and the cytosolic protein FabI was used as a negative control. The positive control used was the *L. pneumophila* effector RalF. Both controls, in addition to wild-type and mutant HipT_Lp_(D197Q, D219Q), were expressed from the pXDC61 vector (induced with 500 μM IPTG) as a fusion with the TEM-1 protein and infections were performed with the indicated multiplicities of infection (MOIs). Quantified fluorescence is reported as the ratio of blue fluorescence (translocation) to green fluorescence (no translocation). The bar chart shows the mean ± standard deviation of 3 biological replicates and is representative of 3 independent experiments. The dashed grey line indicates a blue/green fluorescence ratio of 1, which was used as a threshold for translocation. **(B)** Expression of HipT_Lp_ in *S. cerevisia*e BY4742 cells. Yeast cultures carrying HipT_Lp_ cloned into the pAG425GAL expression vector, or a vector only control, were grown overnight and spotted in serial dilutions onto media containing either 2% glucose (repression) or 2% galactose (expression) and grown for 2 days. Spotting assays were performed in biological triplicate and images are representative of 3 independent experiments.

### HipBST_Lp_ is a functional tripartite toxin-antitoxin system

We next wondered whether the HipBST_Lp_ module could serve as a functional TA system. To test this, we cloned *hipT*_Lp_ into an inducible plasmid and examined the impact of its expression on bacterial growth. HipT_Lp_ expression inhibited growth in both an *L. pneumophila* strain with the endogenous locus deleted (Δ*hipBST*) and *E. coli* (Figure 3A, S3A). This effect was neutralized in *L. pneumophila* by co-expression of HipS_Lp_, but not HipB_Lp_ (Figure 3B, S3B). No growth inhibitory effects were observed for expression of HipB_Lp_ or HipS_Lp_, either alone or in combination (Figure S3C). These results are consistent with the reported functionality of the HipBST_Ec_ system (12) and confirm that HipBST_Lp_ is a functional TA module.

**Figure 3.**
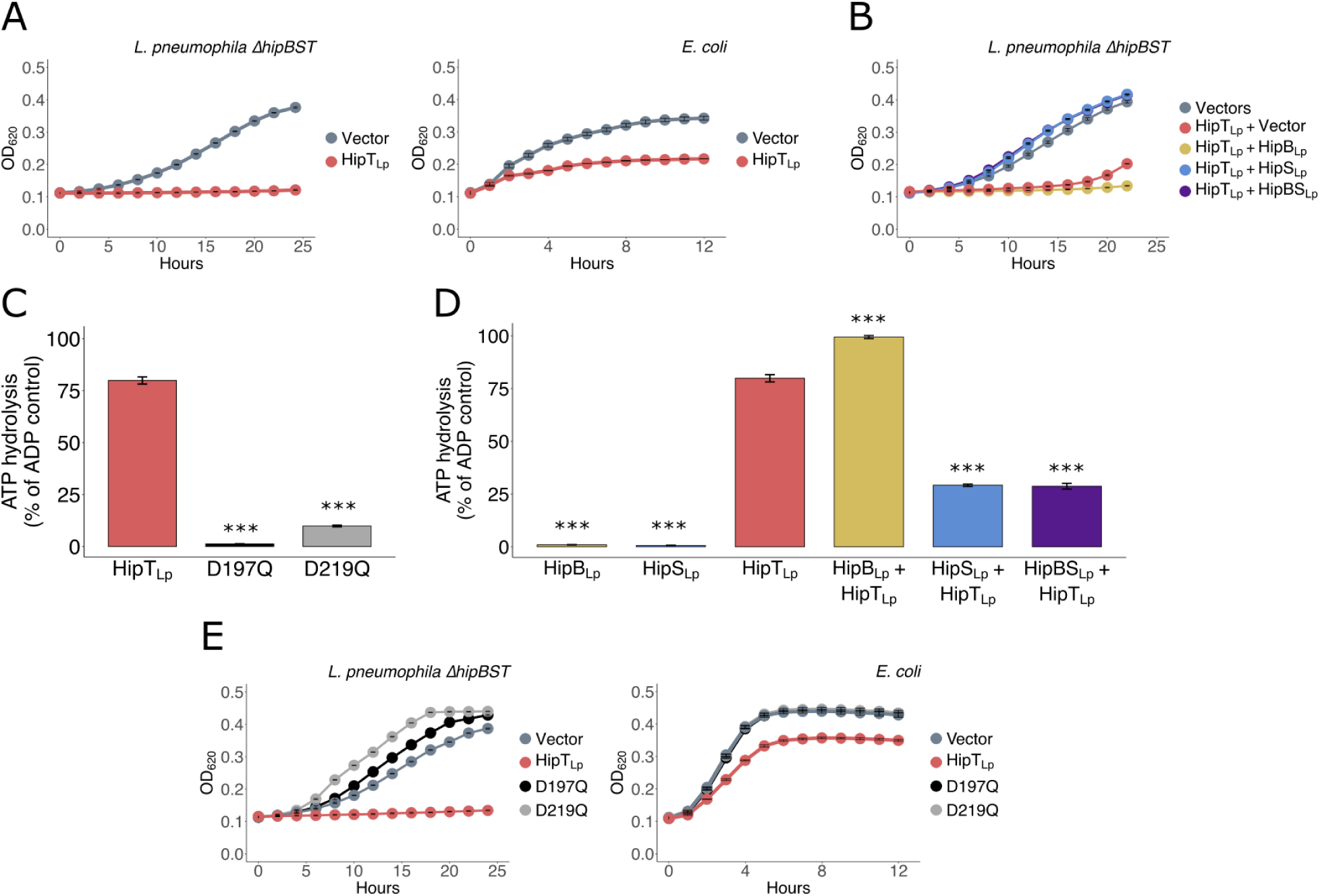
HipBST_Lp_ is a functional tripartite TA system that restricts growth via HipT_Lp_ kinase activity. **(A)** Expression of HipT_Lp_ in *L. pneumophila* cells with the endogenous *hipBST* locus deleted (Δ*hipBST*) and *E. coli* (TOP10) cells. Expression was induced with IPTG (100 μM) for the pJB1806 expression vector used in *L. pneumophila* and arabinose (0.2%) for the pBAD18 vector used in *E. coli*. **(B)** HipT_Lp_ co-expression with HipB_Lp_ and HipS_Lp_ in *L. pneumophila* Δ*hipBST* cells. HipT_Lp_ was expressed from the pJB1806 vector and HipB_Lp_, HipS_Lp_ were co-expressed from the pNT562 vector. Expression was induced with IPTG (100 μM). **(C)** ADP-Glo kinase assay with purified recombinant His_6_-SBP-tagged HipT_Lp_. Both wild-type HipT_Lp_ and HipT_Lp_ with substitutions in two conserved catalytic residues (D197Q, D219Q) were assayed. Reactions contained 1 μg of protein and were incubated at 37°C for 30 min. **(D)** ADP-Glo kinase assay with purified recombinant His_6_-SBP-tagged HipB_Lp_, HipS_Lp_, and HipT_Lp_. Reactions contained 1 μg of each protein and were incubated at 37°C for 30 min. **(E)** Expression of HipT_Lp_ with mutations in two conserved catalytic residues (D197Q, D219Q) in *L. pneumophila* Δ*hipBST* and *E. coli* (TOP10) cells. Expression was induced with IPTG (100 μM) for the pJB1806 expression vector used in *L. pneumophila* and arabinose (0.2%) for the pBAD18 vector used in *E. coli*. All growth curves show the mean ± the standard deviation of 3 biological replicates. Data are representative of 3 independent experiments. All kinase assays show the mean ± the standard deviation of 3 technical replicates. Data are representative of a minimum of 2 independent experiments. Statistical hypothesis testing for the kinase assays was performed with a two-tailed Student’s *t*-test and each sample was compared to HipT_Lp_. ***=*p*-value<0.0001; α (0.05) was Bonferroni corrected for multiple hypothesis testing.

HipT_Ec_ has been shown to be a kinase and this activity is required for its toxicity to the cell (12). We therefore tested whether HipT_Lp_ also displayed kinase activity. We purified wild-type HipT_Lp_, HipT_Lp_(D197Q), and HipT_Lp_(D219Q) (Figure S3D), and tested for ATP hydrolytic activity using the ADP-Glo kinase assay. Wild-type HipT_Lp_ demonstrated substantial activity, whereas the activity of both mutants was abrogated (Figure 3C). ATP hydrolysis was also reduced for wild-type HipT_Lp_ by the addition of the kinase inhibitor FSBA (Figure S3E). Coincubation of purified recombinant HipS_Lp_ (Figure S3D) with HipT_Lp_ neutralized activity *in vitro* (Figure 3D), whereas coincubation with recombinant HipB_Lp_ (Figure S3D) did not, and instead an increase in activity was observed with this combination. The cause of this is unclear, though it may be a consequence of promiscuous phosphorylation of HipB_Lp_ by HipT_Lp_. The D197Q and D219Q substitutions also eliminated the growth inhibitory phenotype of HipT_Lp_ in the Δ*hipBST* strain and *E. coli* (Figure 3E, S3F-G), confirming that kinase activity is required for cellular toxicity. Taken together, these results demonstrate that the HipBST_Lp_ locus encodes a functional TA system, in which HipT_Lp_ toxicity results from its kinase activity and neutralization is performed by HipS_Lp_ rather than HipB_Lp_.

### The HipBST_Lp_ system has the capacity for complex regulatory dynamics

Antitoxins of type II TA systems, such as HipB in the HipBA system, neutralize their cognate toxins via direct physical interaction (11). As HipS_Lp_ functions as the antitoxin in the HipBST_Lp_ system, we wondered if it physically interacts with the HipT_Lp_ toxin. To test this, we examined the binary protein-protein interactions of the HipBST_Lp_ system using the yeast two-hybrid assay. Our results showed that HipT_Lp_ could physically interact with both HipB_Lp_ and HipS_Lp_ independently, but these two proteins could not themselves physically interact (Figure 4A, S4A). To test whether all three proteins could stably associate, we repeated the experiment with the addition of constitutively expressed HipT_Lp_. In this setup, HipB_Lp_ and HipS_Lp_ came in close enough proximity to produce a detectable interaction (Figure 4A, S4A), suggesting that HipT_Lp_ could interact with both proteins simultaneously and serve as a scaffold for a tripartite complex.

**Figure 4.**
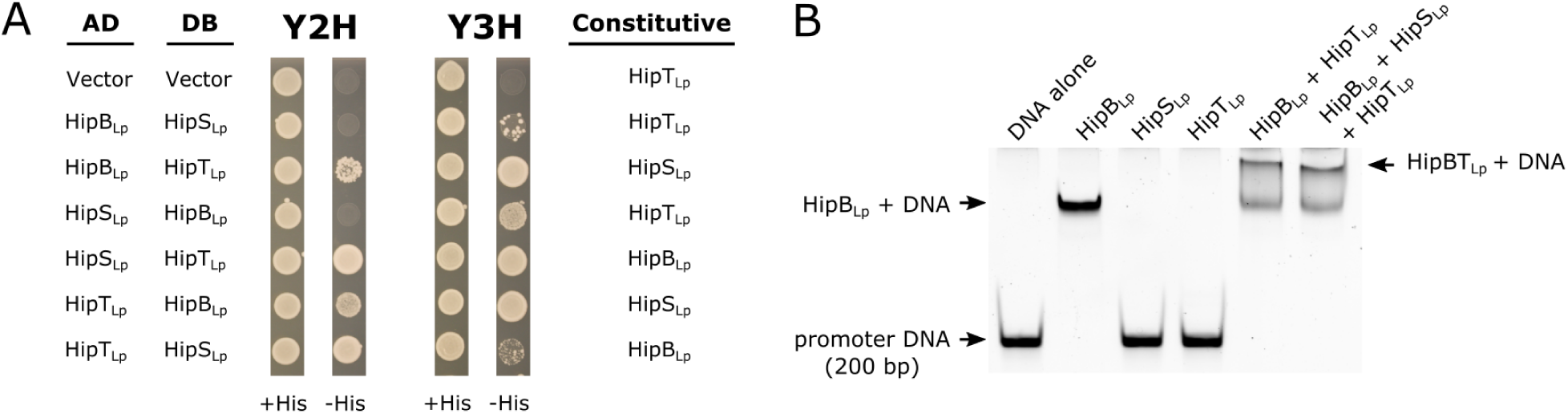
The HipBST_Lp_ system has the capacity for complex binary and ternary regulatory dynamics. **(A)** Yeast two-hybrid (Y2H) experiments testing for binary physical interactions in the HipBST_Lp_ tripartite proteins. Genes cloned into the pDEST-AD and pDEST-DB Y2H vectors are indicated, and representative images are shown of *S. cerevisia*e Y8800 growth in the presence (+His) and absence (-His) of histidine. Growth in the absence of histidine can only occur through a stable protein-protein interaction, due to reconstitution of the GAL4 transcription factor (AD and DB domains) and subsequent expression of the *HIS3* reporter gene downstream of the *GAL1* promoter. Yeast two-hybrid experiments were also performed with a third protein constitutively expressed from the pAG416 vector (Y3H). Genes cloned into pAG416 are indicated. **(B)** Electrophoretic mobility shift assay (EMSA) performed with recombinant purified His_6_-SBP-tagged HipBST_Lp_ proteins (all at 0.2 μM) and a 200 bp DNA fragment encompassing the region upstream (promoter) of the *hipBST_Lp_* locus. The gel was stained with SYBR Green and protein-DNA complexes are indicated.

In the HipBA system, HipB both neutralizes HipA and regulates transcription of the *hipBA* locus by binding to the upstream promoter region (57). We next asked whether toxin neutralization and autoregulation of *hipBST*_Lp_ transcription are decoupled, given the tripartite architecture of the HipBST_Lp_ system. We tested all three proteins for binding of 200 bp upstream (*hipBST_Lp_* promoter) sequence using an electrophoretic mobility shift assay (EMSA). Co-incubation with HipB_Lp_ slowed the migration of the DNA fragment (Figure 4B), whereas HipS_Lp_ and HipT_Lp_ did not, suggesting that the transcriptional regulation of the *hipBST_Lp_* locus is indeed distinct from toxin neutralization. We also observed a secondary gel shift when HipB_Lp_ was incubated with HipT_Lp_, either alone or with HipS_Lp_ also present, though it was unclear whether HipS_Lp_ was in fact interacting with HipT_Lp_ in this reaction. Additionally, the affinity of HipB_Lp_ for promoter DNA appeared to slightly increase in the presence of HipT_Lp_ (Figure S4B). Taken together, these results reveal the different binary and ternary interactions that can occur in the HipBST_Lp_ system, and suggest that transcriptional regulation by HipB_Lp_ is independent of HipT_Lp_ neutralization. HipB_Ec_ has also been reported to regulate transcription of the *hipBST*_Ec_ locus (58), however our data demonstrate the occurrence of a HipB_Lp_-HipT_Lp_-DNA complex, suggesting that interaction dynamics between ternary protein combinations may serve as additional layers in a larger regulatory program.

### HipBST_Lp_ neutralization exploits the P-loop ejection mechanism of HipA_Ec_

To understand the structural basis for neutralization in the HipBST_Lp_ system, we solved the *apo* structure of HipB_Lp_ and co-crystal structure of HipS_Lp_-HipT_Lp_ (Table S7). HipB_Lp_ is highly similar to HipB_Ec_ (RMSD 1.6-2.2 Å over approximately 70 Cɑ atoms) and adopts a dimeric helix-turn-helix conformation, consistent with its capacity to bind DNA (Figure 5A). Strikingly, the structural conformation of the HipS_Lp_-HipT_Lp_ complex is nearly identical to HipA_Ec_ (RMSD 2.8 Å over 421 Cɑ atoms, when considering HipS_Lp_ and HipT_Lp_ as a single chain) (Figure 5B), despite the functional divergence between the systems. Given that HipS_Lp_ serves the role of antitoxin, we wondered how neutralization could be achieved in this orientation. To address this, we compared the HipS_Lp_-HipT_Lp_ structure to previously solved HipA_Ec_ structures. Independent of neutralization by HipB_Ec_, HipA_Ec_ is capable of self-inhibition through intermolecular phosphorylation of its S150 residue (59). This site is found within the catalytic P-loop, and its phosphorylation or mutation to alanine yields a conformational shift whereby the P-loop becomes ejected and solvent exposed, thereby inactivating the toxin (Figure 5C). Part of the P-loop in our HipS_Lp_-HipT_Lp_ structure was unresolved—suggesting it was in a flexible or dynamic state—but the resolved portion (residues 50-56) was oriented in a manner highly similar to the ejected and inactive HipA_Ec_ P-loop (Figure 5C). This suggests that the interaction between HipS_Lp_ and HipT_Lp_ facilitates P-loop ejection to neutralize the toxin, revealing a mode of action that exploits an intrinsic property of HipA. During our analyses, the structures of both HipT_Lp_ and HipS_Lp_-HipT_Lp_ were reported by another group (21). These illustrate the orientation of the P-loop in its internalized and catalytically active state, and as in our HipS_Lp_-HipT_Lp_ structure, this motif becomes unresolved and likely ejected upon HipS_Lp_ binding. These findings are therefore consistent with ours and similar structural observations have also been reported for the HipBST_Ec_ system (58), suggesting a conserved P-loop ejection model of HipBST neutralization.

**Figure 5.**
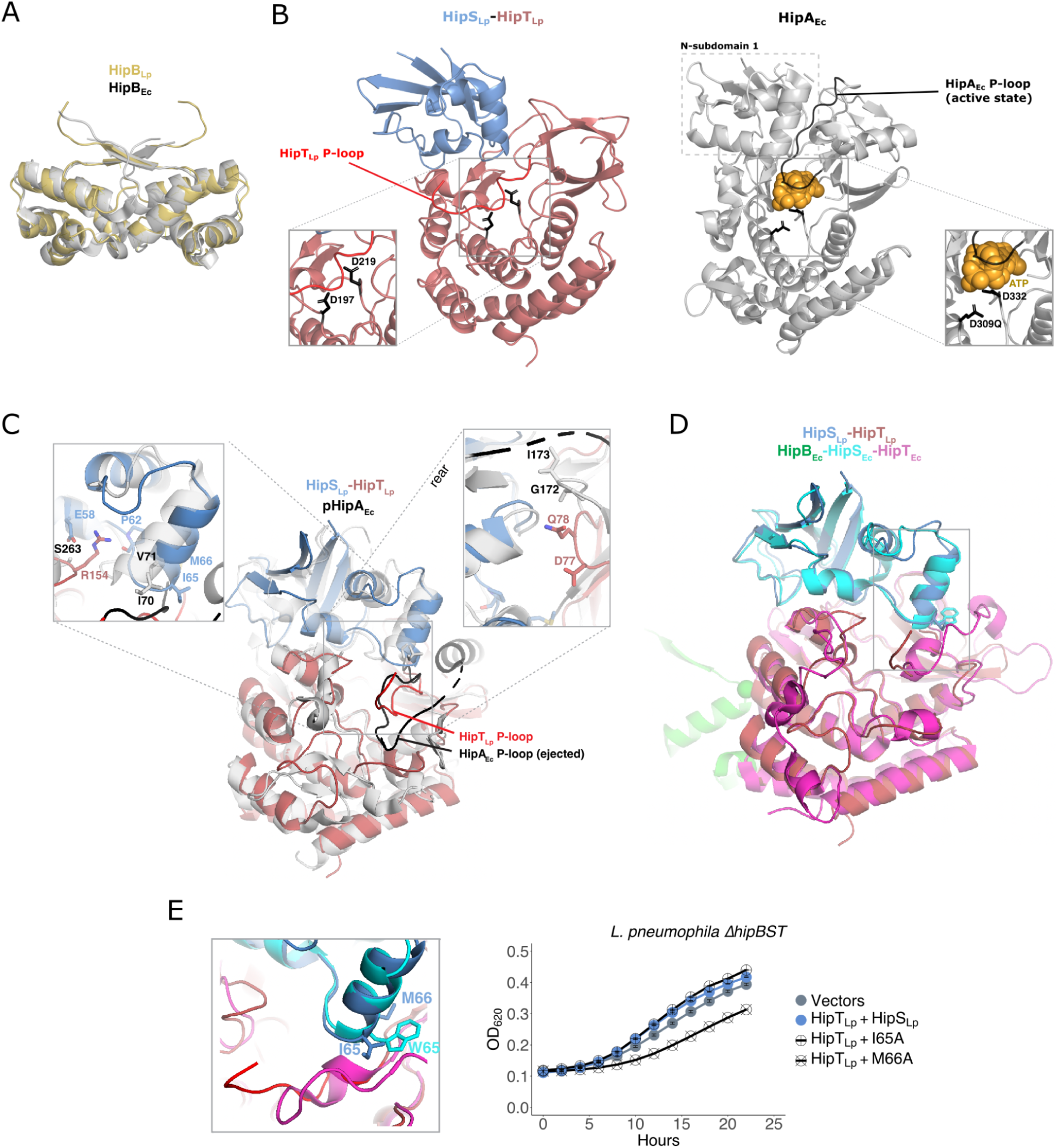
HipT_Lp_ neutralization by HipS_Lp_ utilizes a P-loop ejection mechanism. **(A)** Alignment of the HipB_Lp_ structure with HipB_Ec_ (4YG1). **(B)** Comparison of the HipS_Lp_-HipT_Lp_ co-crystal structure with HipA_Ec_ (3DNT) in which the HipA_Ec_ P-loop motif is in the internalized and active conformation. The HipA_Ec_ N-subdomain 1, corresponding to HipS_Lp_, is indicated with a dashed box. The P-loops of both HipT_Lp_ (red) and HipA_Ec_ (black) are indicated. ATP in the HipA_Ec_ catalytic pocket is depicted with yellow spheres. Insets display the catalytic pocket of both kinases, along with 2 conserved residues involved in ATP coordination (D197, D309Q) and Mg^2+^ binding (D219, D332). **(C)** Alignment of the HipS_Lp_-HipT_Lp_ structure and pHipA_Ec_ (3TPE), in which the P-loop motif is autophosphorylated and in the inactive, ejected conformation. Insets display two regions of variation between the HipS_Lp_-HipT_Lp_ interaction interface and HipA_Ec_: a shifted helix in HipS_Lp_ (inset on the left) and a loop in the N-terminal region of HipT_Lp_ (inset on the right). Key residues in each region are labelled. **(D)** Alignment of the HipS_Lp_-HipT_Lp_ structure with the structure of HipBST_Ec_ (7AB5). **(E)** Left: inset from **(F)** comparing the tryptophan residue in HipS_Ec_ (W65) with the methionine (M66) and isoleucine (I65) residues at the same position in HipS_Lp_. Right: Co-expression of HipT_Lp_ (pJB1806) with wild-type or mutant (I65A, M66a) HipS_Lp_ (pNT562) in *L. pneumophila* Δ*hipBST* cells. Expression was induced with IPTG (100 μM). Growth curves show the mean ± the standard deviation of 3 biological replicates. Data are representative of 3 independent experiments.

To better understand the evolution of this unique mechanism, we sought to identify sites of divergence between the HipS_Lp_-HipT_Lp_ complex and HipA_Ec_. Two major conformational changes are present in HipT_Lp_ loop 75-80 and HipS_Lp_ helix 65-66, relative to the equivalent regions in HipA_Ec_ (171-172 and 70-71), which shift these regions inward and impinge on the space where the P-loop would be internalized in its active state (Figure 5C). We identified two residues in HipT_Lp_ (D77, Q78) that occupy this space and are not conserved in HipA_Ec_ (Figure S1A). Substitution of either residue with alanine did not impair HipT_Lp_ toxicity (Figure S5A), however it also did not prevent neutralization by HipS_Lp_ (Figure S5B). The HipT_Lp_ residue R154 is also not conserved in HipA_Ec_ or HipT_Ec_ and appears to contribute to the interaction between toxin and antitoxin through the formation of a hydrogen bond to the backbone of HipS_Lp_(P62) and a salt bridge with HipS_Lp_(E58) (Figure 5C). However, mutation of this residue to alanine (R154A) was again not sufficient to impair the neutralization of HipT_Lp_ by HipS_Lp_ (Figure S5A-B).

### HipT_Lp_ neutralization by HipS_Lp_ does not rely on the tryptophan residue utilized by HipS_Ec_

We next compared the *L. pneumophila* and *E. coli* HipBST systems directly, as a structure of HipBST_Ec_ was recently reported (58) (Figure 5D). In the HipBST_Ec_ system, neutralization has been reported to depend on a tryptophan residue in HipS_Ec_ (W65) that projects into the P-loop containing pocket of HipT_Ec_ (Figure 5D-E). This residue appears critical for HipS_Ec_ function and is conserved across numerous HipS_Ec_ homologs, but is absent from HipS_Lp_. Instead, this site is occupied by much smaller methionine and isoleucine residues (I65, M66). We substituted both residues with alanine and observed a partial reduction in HipT_Lp_ neutralization by HipS_Lp_(M66A), but no effect for HipS_Lp_(I65A) (Figure 5E, S5C).

Consistent with these results, neither substitution impaired the physical interaction between HipS_Lp_ and HipT_Lp_ (Figure S5D). In summary, the HipS_Lp_-HipT_Lp_ P-loop ejection mechanism is broadly conserved with HipBST_Ec_, yet is achieved through different motifs and interactions within the toxin-antitoxin interface.

### HipT_Lp_ toxicity is not inhibited by autophosphorylation of its P-loop serine residue

The HipBST_Ec_ system was recently shown to contain a double serine motif in the HipT_Ec_ P-loop (S^57^IS^59^) that is proposed to allow for a dual autoregulatory dynamic in that system (58). This motif is absent from HipT_Lp_, which contains only a single serine in this position (S54), similar to HipA_Ec_ (S150) (Figure S1A). While many of the previously reported HipT homologs from the Gammaproteobacteria (51) contain either an SxS motif, or some combination of double S/T residues, the corresponding motif in HipT_Lp_ (SVQ) is absent in these homologs but is either conserved or nearly identical (SIQ) across the HipT_Lp_ homologs in *Legionella* species. It has been shown previously that autophosphorylation or substitution of the single P-loop serine in HipA_Ec_ leads to P-loop ejection and loss of activity (55, 59). We therefore wondered what consequence modifying this residue would have on HipT_Lp_ toxicity. To test this, we constructed mutations that both ablate (S54A) and mimic (S54D) phosphorylation at this position. Neither mutation inhibited toxicity (Figure S5E), which was surprising given that the corresponding mutation in HipT_Ec_ (S57A) renders the protein non-toxic (58), as does the S150A mutation in HipA_Ec_ ^(^55^)^, and autophosphorylation of HipA_Ec_ renders it unable to bind ATP and retain catalytic activity (59). These findings are supported by recent structural work demonstrating that HipT_Lp_ can bind ATP despite being autophosphorylated (21), and highlight an unusual autoregulatory difference between these systems.

### The cellular target of HipT_Lp_ is different from those of characterized HipT and HipA toxins

Despite their phylogenetic divergence, the HipBST_Lp_ system functions similarly to HipBST_Ec_. Given the dearth of comparisons between TA homologs across distantly related bacteria, we chose to further explore the functional conservation between systems by testing whether they modified the same substrate. HipT_Ec_ was previously shown to phosphorylate tryptophan tRNA-ligase (TrpS) in order to arrest cellular growth (12), whereas the canonical target of HipA_Ec_ is glutamyl tRNA-synthetase (GltX) (60, 61). TrpS and lysine tRNA-ligase (LysS) have also been reported as substrates for the HipBA systems in *Caulobacter crescentus* (62, 63), however the primary targets of these systems remain uncertain. In *E. coli*, co-expression of TrpS or GltX with HipT_Ec_ or HipA_Ec_, respectively, rescues the growth inhibitory effect of the toxins. In fact, *E. coli* TrpS was sufficient to rescue the toxicity of HipT homologs from *Haemophilus influenzae* and *Tolumonas auensis* as well (12). We therefore tested whether overexpression of TrpS_Lp_ or GltX_Lp_ could rescue HipT_Lp_-induced growth inhibition when co-expressed in *L. pneumophila*. Surprisingly, neither of these proteins could alleviate HipT_Lp_ toxicity (Figure 6A, S6A), suggesting that HipT_Lp_ poisons the cell by targeting one or more previously undescribed substrate(s).

**Figure 6.**
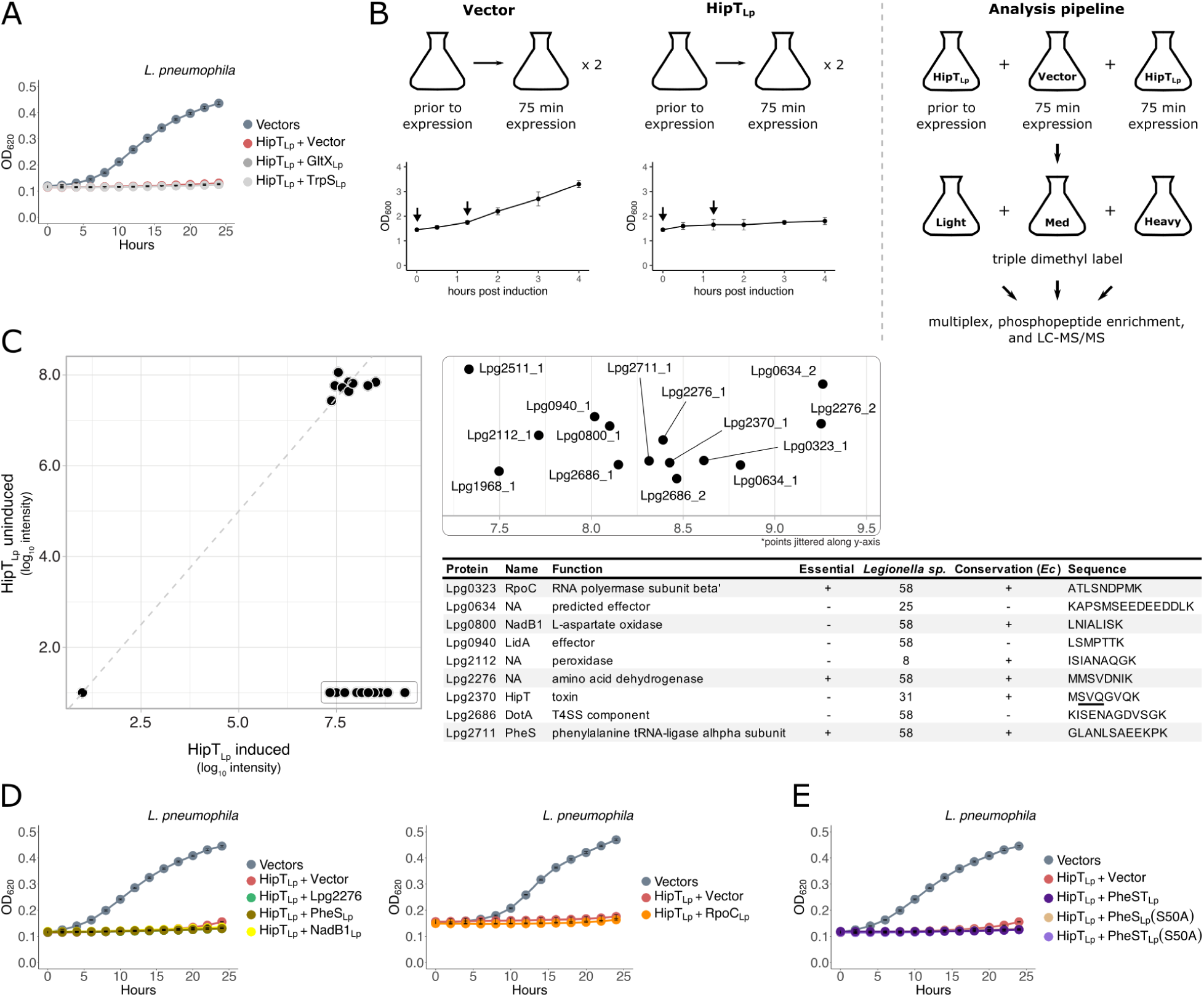
HipT_Lp_ has an unknown cellular target that is not conserved with characterized HipT or HipA toxins. **(A)** Co-expression of HipT_Lp_ (pJB1806) with GltX_Lp_ or TrpS_Lp_ (pNT562) in *L. pneumophila* cells. **(B)** Overview of phosphoproteomic experimental design. Left: cultures of *L. pneumophila* Δ*hipBST* carrying either pJB1806 or pJB1806::*hipT_Lp_* were grown to mid-log phase in the presence of 0.5% glucose, after which expression was induced by the addition of IPTG (100 μM). Prior to and 75 mins post induction (indicated by arrows on growth curves), cells were harvested for phosphoproteomic analysis. This experiment was performed independently twice and growth curves show the mean ± the standard deviation from the 2 independent experiments. Right: cell lysates from cultures expressing HipT_Lp_ at both time points, and the vector control post induction, were dimethyl labelled and multiplexed, enriched for phosphopeptides, and analyzed by mass spectrometry. The HipT_Lp_ uninduced, vector induced, and HipT_Lp_ induced samples were labelled with the light, intermediate (med), and heavy channels, respectively. **(C)** Phosphopeptides detected in both HipT_Lp_ uninduced and induced samples (left), and those enriched during HipT_Lp_ expression only (inset), are displayed. The dashed grey line is y=x for comparison between channels. In the inset, the suffix denotes the peptide number seen from a given ORF. The intermediate (vector induced) channel is excluded from this plot for simplicity and because the results did not change the list of candidate substrates. The table contains candidate phosphoproteins that were observed across both replicates (excludes Lpg1968 and Lpg2511) along with their function, essentiality in broth, conservation in 58 *Legionella* species and *E. coli* (*Ec)*, and detected phosphopeptide sequence. Candidates chosen for subsequent validation are highlighted in grey. The HipT_Lp_ phosphosite is underlined. **(D)** *L. pneumophila* cells co-expressing HipT_Lp_ (pJB1806) with phosphoproteomic candidates (pNT562) that were either essential, or highly conserved in both *Legionella* and conserved in *E. coli*. **(E)** Co-expression of HipT_Lp_ with PheST_Lp_ or phosphomimetic mutants of PheS_Lp_(S50A) in *L. pneumophila* cells. Growth curves show the mean ± the standard deviation of 3 biological replicates. Expression was induced using IPTG (100 μM) for all constructs. Data are representative of 3 independent experiments.

GltX and TrpS were originally identified as the targets of HipA_Ec_ and HipT_Ec_ by screening *E. coli* genomic libraries for genes whose overexpression rescued toxin-induced growth inhibition (12, 60) or cold sensitivity (61). Given the observed toxicity of HipT_Lp_ in *E. coli* cells, we used the same approach to search for putative toxin targets. We pooled and transformed the *E. coli* ASKA library into BL21 cells expressing HipT_Lp_ (pCDF1-b) and screened the resulting transformants. We observed robust growth inhibition upon HipT_Lp_ expression using an empty vector control (pCA24N), further demonstrating the toxicity of HipT_Lp_ in *E. coli* (Figure S6B). When cells were transformed with the ASKA library pool, we recovered a small number of transformants across multiple screen replicates (Figure S6B). Despite numerous clones exhibiting a stable growth rescue phenotype, sequencing revealed no clear enrichment of any genes or pathways across experimental replicates and subsequent attempts at validation in *L. pneumophila* were unsuccessful.

As rescue screening of the ASKA library did not produce any strong candidate HipT_Lp_ targets, we next pursued an orthogonal approach that was not reliant on growth rescue for substrate identification. Phosphoproteomic analysis was used previously to identify TrpS as a substrate of *C. crescentus* HipA2 (63) and to demonstrate the enrichment of GltX phosphorylation in the HipA_Ec_ phosphoproteome (64). We therefore performed a phosphoproteomic screen in *L. pneumophila* to detect substrates that were only modified under conditions of HipT_Lp_ overexpression, relative to both uninduced HipT_Lp_ and an empty vector control (Figure 6B). From this, we identified a small number of proteins that were enriched for phosphopeptides during HipT_Lp_-induced growth inhibition across two biological replicates (Figure 6C). We also observed that HipT_Lp_ is phosphorylated on S54, thereby demonstrating the occurrence of this modification *in vivo*. Of the 8 candidate substrates identified, 3 are known to be essential for growth (17, 18), 6 are conserved across all *Legionella* species, and 5 have orthologs in *E. coli* (Figure 6C). While HipA_Ec_ has been shown to phosphorylate a large pool of proteins, only co-expression with GltX is sufficient to rescue its growth inhibitory phenotype (64). In order to validate our candidate substrates, we cloned a subset of hits that were either essential or highly conserved and tested them for growth rescue when co-expressed with HipT_Lp_. Under these conditions, no putative substrate was able to rescue growth inhibition in *L. pneumophila* (Figure 6D, S6C).

One of the hits we detected was the alpha subunit of phenylalanine tRNA-ligase (PheS). Given that both GltX and TrpS are tRNA-ligases, this class of target would be consistent with other Hip toxins. Interestingly, phenylalanine tRNA-ligase is composed of two separate subunits, whereas GltX and TrpS are both single proteins. To test whether the complete phenylalanine tRNA-ligase (PheST_Lp_) was required for growth rescue, we co-expressed it with HipT_Lp_ in *L. pneumophila*. However, we did not observe any rescue of growth inhibition (Figure 6E, S6D). From our phosphoproteomic data, we also observed that PheS was phosphorylated on the S50 residue during HipT_Lp_ overexpression. We next tested whether mutation of this site to ablate phosphorylation (S50A) would prevent growth inhibition, however neither PheS(S50A) nor PheST(S50A) were able to rescue *L. pneumophila* growth when co-expressed with HipT_Lp_ (Figure 6E, S6D). While these proven approaches were unable to identify a single cellular target of HipT_Lp_ responsible for the observed growth defect, they nevertheless suggest that the cellular target of HipT_Lp_ is likely not conserved with HipT_Ec_ or any characterized HipA homologs.

## DISCUSSION

The ubiquity of toxin-antitoxin systems in bacterial genomes and their capacity for selfish maintenance have provided strong evidence for their role as parasitic elements. Broader functionality of chromosomal TA systems and the conservation of system function across different bacterial hosts remain open questions (7). Answering these questions is challenging, due in part to the poor conservation of many modules even across closely related strains and species. Given the plastic nature of the accessory genome and the patchy distribution of TA systems, the occurrence of highly conserved elements would suggest selection beyond mere addiction. Here we characterize a HipBST TA system in *L. pneumophila* that is highly conserved across *Legionella* species, despite the genomic divergence within this genus. Indeed, more than half of the *Legionella* species we searched have either a complete HipBST module (28/58 species) or individual HipBST genes (8/58 species). This level of conservation is notable, in that it is higher than both the average accessory gene (26/58 species) and *L. pneumophila* effector (16/58).

In characterizing HipBST_Lp_, we detected a large number of distant homologs across diverse bacterial taxa. This greatly expands the sequence space of HipBST systems beyond the 48 previously reported (51) and reveals considerable diversity within this newly described TA family. We were surprised to discover that HipBST_Lp_ and its homologs form a distinct subclade from HipBST_Ec_. While HipBST_Ec_ homologs are predominantly found in the Gammaproteobacteria class, homologs of the HipBST_Lp_ system are almost entirely restricted to the *Legionella* genus within that class but are widely distributed in other taxonomic groups, such as the FCB superphylum. This could suggest a shared environmental niche, common functional role, or frequent DNA exchange between these taxa and *Legionella* species. However, it is unclear why this divergent HipBST variant is so prevalent in *Legionella* relative to the rest of the taxa in the Gammaproteobacteria. Importantly, previous work has only compared HipBST systems that are closely related to HipBST_Ec_ (12). As the sequence space of HipBST_Lp_ homologs appears to be both more diverse and taxonomically distributed (Figure 1C-D), this offers the potential to reveal even more breadth of TA biology within a single family.

During our genomic search, we discovered a putative duplicate HipBST system downstream of HipBST_Lp_. This locus (*lpg2377-80*) encodes 5 predicted ORFs and is only found in the *L. pneumophila* genome (Figure S7A). Its unusual architecture is achieved through the further splitting of both *hipS* and *hipT* into two ORFs each, resulting in the separation of the P-loop and kinase core in HipT. Interestingly, all catalytically critical motifs remain conserved in split-HipT (Figure S7A). The overlaps between the split-*hipS* and split-*hipT* ORFs are 23 and 41 bp respectively, which is larger than the 4 bp overlap observed in all other ORF boundaries in both systems. As large ORF overlaps can impact translational coupling in bacterial operons (65), this may indicate significant regulatory evolution in this system. To determine whether this system could function as a TA system, we expressed both split-HipT_Lp_ and split-HipS_Lp_ in *L. pneumophila* cells. Despite the preservation of the P-loop motif, the expression of either split protein pair, alone or in combination, did not exhibit a growth inhibitory effect (Figure S7B). It thus remains unclear why this locus has evolved its unusual genetic architecture and has been retained in the genome. We also observed a different split-*hipT* organization in the HipBST system encoded by *Legionella gormanii*, where *hipT* has been split in a similar manner to *lpg2379-80,* but in this case the upstream ORF is truncated due to a frameshift and the catalytic P-loop is no longer intact (Figure S7C). Thus this system appears to be in a state of decay or functional divergence. Importantly, while it remains to be seen how these novel split-protein genetic architectures affect system functionality, their growing diversity attests to the substantial modularity and evolvability of the Hip TA systems.

A striking feature of the HipBST_Lp_ system is the previous claim that HipT_Lp_ is a *L. pneumophila* effector (20, 21). This would represent an exceedingly rare example of toxin-antitoxin system/eukaryotic effector bifunctionality, as there is limited evidence of TA modules being repurposed as interdomain translocated substrates (7). However, we found no evidence of HipT_Lp_ translocation beyond levels of the negative control FabI. Instead, the broad taxonomic distribution of HipBST_Lp_ homologs we observed likely extends to many species that do not deliver effectors into host cells, further arguing against this functionality. HipT_Lp_ was first hypothesized to be an effector due to the presence of a putative C-terminal translocation motif (66), however subsequent re-examination determined it was unlikely to encode a true secretion signal (67). Our inability to observe HipT_Lp_ translocation, despite similar methodology, argues for caution in ascribing functionality to HipT_Lp_ within the eukaryotic host. Indeed, it should be noted that in previous work, HipT_Lp_ was reported to be an effector based solely on a single qualitative micrograph demonstrating exceedingly low translocation efficiency (20, 21). Given that cytosolic proteins (such as FabI) can be translocated at low frequencies by the T4SS (29, 68), basal level of secretion should not be mistaken for functional translocation and warrants confirmation with orthologous methodologies. In the absence of this, exceeding a threshold ratio of quantified fluorescence should be the necessary standard for effector validation (29, 30).

In this work, we demonstrate that HipBST_Lp_ is a TA system—the core functionality of which is conserved with HipBST_Ec_—and provide a detailed comparison of the same TA family across different bacterial species. A recent report on HipBST_Lp_ ^(^21^)^ included the puzzling claim that HipT_Lp_ does not inhibit growth in *E. coli*. We routinely observed that overexpression of HipT_Lp_ consistently inhibits *E. coli* growth, using multiple strains, vectors, and induction methods (Figure 3A, S3F-G, S6B). Our detection of a physical interaction between HipB_Lp_ and HipT_Lp_, by both yeast two-hybrid and EMSA assays, is also contradictory to the previous report (21) which found no such interaction. The absence of an interaction between HipB_Lp_ and HipT_Lp_ would suggest a regulatory dynamic wherein only neutralized HipT_Lp_ would alter the regulatory activity of HipB_Lp_. Instead, our observed interaction between HipB_Lp_ and HipT_Lp_ raises the possibility of multi-layered regulation in the HipBST_Lp_ system, whereby both binary and ternary protein interactions could influence system expression levels in an expanded form of conditional cooperativity (Figure 7).

**Figure 7.**
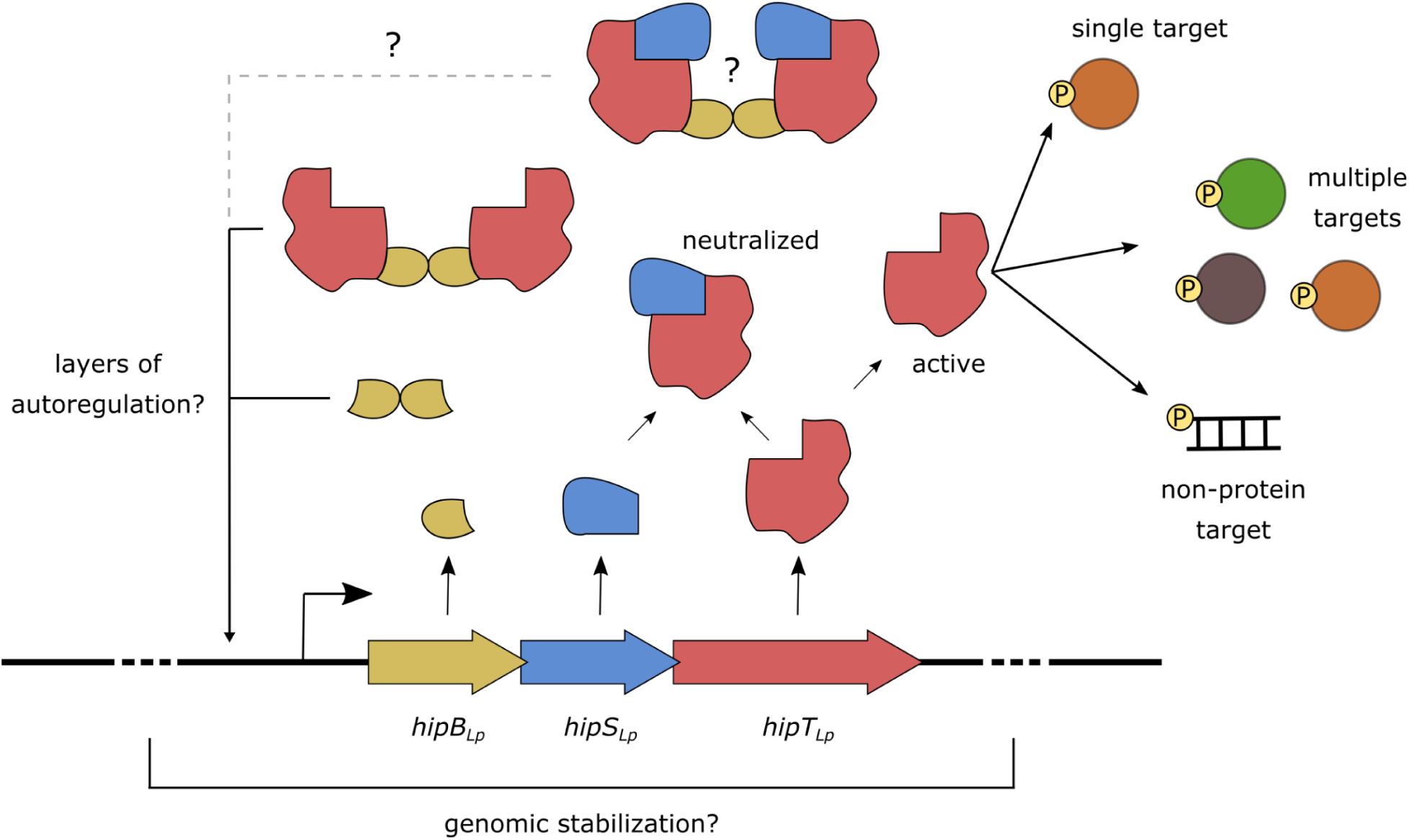
Model of the HipBST_Lp_ toxin-antitotin system. HipT_Lp_ is neutralized by physical interaction with HipS_Lp_ and HipB_Lp_-HipS_Lp_-HipT_Lp_ can stably associate, however it remains uncertain whether the ternary complex forms a hexamer through HipB_Lp_ dimerization. HipB_Lp_ and HipB_Lp_-HipT_LP_ can bind promoter DNA to regulate *hipBST_Lp_* expression, creating the possibility for two layers of transcriptional regulation in the system. It is unknown whether HipB_Lp_-HipS_Lp_-HipT_Lp_ can also bind DNA, thus adding another layer to the regulatory program. The target of HipT_Lp_ remains unknown, but is not conserved with characterized HIpT or HipA toxins. It could therefore be an entirely new protein target, multiple proteins in the cell, or a non-protein cellular target. HipBST_Lp_ homologs are commonly found in accessory genomic clusters of non-essential genes across *Legionella* species genomes. One hypothesis is that these systems serve to stabilize these highly plastic regions of the genome, thereby maintaining the rich diversity of effectors found in *Legionella* species.

We were surprised to find that the HipS_Lp_-HipT_Lp_ complex adopts a conformation that is almost unchanged from that of HipA_Ec_, consistent with recently reported structures for both HipBST_Ec_ ^(^58^)^ and HipBST_Lp_ ^(^21^)^. Neutralization in both HipBST systems is achieved by exploiting a unique property of HipA biology—namely its P-loop ejection mechanism of autoregulation. This represents a fascinating case of functional evolution while maintaining structural congruence. The HipBST_Ec_ and HipBST_Lp_ systems both achieve this functionality through shared structural motifs, while also displaying several key differences, including the HipT_Lp_(R154) and HipS_Lp_(I65,M66) residues. In particular, while the bulky W65 residue in HipS_Ec_ is both critical for toxin neutralization and conserved in many other HipS_Ec_ homologs (58), in HipS_Lp_ this position is occupied by the much smaller I65 and M66 residues. These do not impinge on the catalytic pocket of HipT_Lp_ to the same extent as W65 and are not necessary for HipS_Lp_-HipT_Lp_ physical interaction or neutralization. The absence of an analogous residue to W65 in HipS_Lp_ therefore indicates alternate biochemical means of achieving P-loop ejection may be present across HipBST system homologs.

A double serine motif (S^57^IS^59^) was recently characterized in HipT_Ec_ ^(^58^)^, with the differential autophosphorylation of either serine reported to affect toxin neutralization and activity. This motif is absent from HipT_Lp_, which instead contains a single P-loop serine (S54) similar to the S150 residue in HipA_Ec_. We observed that the phosphorylation state, or substitution of this residue with alanine, does not affect protein toxicity. This was unexpected, given that both pHipA_Ec_ and HipA_Ec_(S150A) cannot bind ATP and are not toxic (59). In support of this unusual behaviour, a structure of pHipT_Lp_ bound to an ATP analogue was recently reported (21), further demonstrating the retention of HipT_Lp_ activity despite its phosphorylation state. Why HipT_Lp_ autoregulation differs in such a critical aspect from HipA_Ec_ is an outstanding question, as this would seemingly prohibit cell detoxification via *trans* autophosphorylation, leading to a kinase with less restricted activity in the cell. Phosphomimetic mutation of either serine residue in HipT_Ec_ (S57D, S59D) does not inhibit toxicity (58), however mutation of either residue to alanine appears to abrogate activity—though conflicting results have been reported (21). Regardless, our findings demonstrate that the autophosphorylation state of HipT_Lp_ is not required for its activity, indicating a shift in the autoregulatory functionality of the HipT_Lp_ kinase.

The most critical difference we observed between the HipBST_Ec_ and HipBST_Lp_ systems is the failure of either GltX_Lp_ or TrpS_Lp_ to rescue HipT_Lp_ toxicity. This was further supported by the absence of both GltX_Lp_ and TrpS_Lp_ in our genomic rescue and phosphoproteomic screens. We did however observe a small set of candidate proteins that were only phosphorylated during HipT_Lp_ overexpression in *L. pneumophila* cells. In particular, we detected the phosphorylation of PheS, which is also a tRNA-ligase like GltX and TrpS, and thus a strong candidate for the target of HipT_Lp_. While none of the putative targets were able to rescue *L. pneumophila* growth when co-expressed with HipT_Lp_, including the complete PheST complex and the phosphoablative mutation in PheS(S50A), this does not eliminate them from consideration. Instead, this may be a consequence of some aspect of *L. pneumophila* or HipT_Lp_ biology that is not conducive to kinase target saturation for growth rescue, or possibly the requirement of multiple co-expressed substrates to overcome toxin activity. In light of our results, it should also be considered whether HipT_Lp_ in fact phosphorylates a protein at all to poison the cell, as the true target may be a nucleic acid or other molecule. Regardless, we report that HipT_Lp_ likely has a different cellular substrate from characterized HipT and HipA toxins, further demonstrating its functional divergence from HipT_Ec_. It will be interesting for future work to assess whether other HipT_Lp_ homologs also have divergent targets from those in the HipT_Ec_ subclade.

Given the conservation of HipBST homologs across *Legionella* species, we wondered whether this system showed any pattern of association with chromosomal subregions or signatures of acquisition. The genome of *L. pneumophila* is organized into discrete clusters of essential and non-essential genes—the latter of which are enriched for effectors and mostly dispensable for broth growth and host infection (17, 18). The HipBST_Lp_ locus is found within one such non-essential cluster in *L. pneumophila* (Figure S8A), and this led us to hypothesize that other HipBST_Lp_ systems might also be found within accessory genomic regions of non-essential genes. To test this, we predicted regions of non-essentiality across the 28 *Legionella* species genomes with HipBST systems. We found that HipBST is frequently associated with non-essential gene clusters (22/28) and that neither the local genetic neighbourhood nor broader cluster composition are well conserved across HipBST_Lp_-containing clusters (Figure S8B). The most common co-occurring genes (25-45% of clusters) were involved in functions such as DNA mobilization and integration, including some in close proximity to the HipBST_Lp_ locus (*lpg2363, lpg2366, lpg2367*). This could therefore represent a common means of HipBST dispersal between *Legionella* species. The high conservation of HipBST modules, contrasted with the poor conservation across clusters, raises the possibility that these loci may serve to stabilize their unique accessory genomic neighbourhood in each species. Indeed, tripartite TA systems are more commonly found on mobile genetic elements and have been hypothesized to function in DNA stabilization (11). In one possible model, integrated HipBST could create a sphere of influence around which genes such as effectors can be stabilized (Figure 7). This would be advantageous for *Legionella* species, which encode vast and diverse effector repertoires—including some with fitness costs in specific hosts (18, 69). Given the broad host range of *Legionella*, maintaining effector diversity likely provides a selective benefit, and similar stabilizing effects have been demonstrated for other chromosomal TA systems (70–73).

Overall, our work provides a detailed characterization of the HipBST TA system in *L. pneumophila*. Given its small and unstudied TA system set, extreme genomic characteristics, and challenging host-associated lifestyle, investigating the TA systems in this species can be a source for new insights into TA biology and function. We find a strong signal of conservation for HipBST within the *Legionella* genus, which is unusual for a TA system and in particular one which is almost completely absent from all other taxa in the Gammaproteobacteria. Despite its frequent occurrence in accessory genomic clusters, this level of conservation suggests that some functionality beyond promiscuous acquisition and addiction is being selected for. We also show that considerable molecular divergence can be found between related or even the same TA system family, which underscores the potential for discovering new biology in the expansive and unexplored sequence space of bacterial genomes. These findings therefore justify caution when generalizing TA system functionality across different bacterial hosts and instead suggest that each system be considered within the context of its specific genomic niche.

## AVAILABILITY

All data have been deposited as a complete submission to the MassIVE repository (https://massive.ucsd.edu/ProteoSAFe/static/massive.jsp) and assigned the accession number (MSV000090735). The dataset is currently available for reviewers at ftp://MSV MSV000090735@<u>massive.ucsd.edu</u>. Please login with username MSV000090735; password: Lin_HipBST_pw. The dataset will be made public upon acceptance of the manuscript.

## ACCESSION NUMBERS

Atomic coordinates have been deposited in the Protein Data Bank with accession codes 8EZR, 8EZS and 8EZT.

## SUPPLEMENTARY DATA

Table S1. Strains used in this study. Table S2. Plasmids used in this study.

Table S3. Oligonucleotides used in this study. Table S4. *Legionella* genome assemblies.

Table S5. HipBA and HipBST homology search results.

Table S5. Phylogeny results for HipBA and HipBST containing species. Table S7. X-ray diffraction data.

Table S8. Phosphoproteomic data.

## ACKNOWLEDGEMENTS

We thank Rosa Di Leo for assistance in cloning the protein purification constructs and Dylan Valleau for help with protein purification for *in vitro* experiments. We thank Cassandra Wong and Brett Larsen for help with the phosphoproteomic methodology and data analysis. We thank Dr. Malene Urbanus and Dr. Beth Nicholson for critical feedback on the manuscript.

## FUNDING

JDL was supported by an Ontario Graduate Scholarship. KTA was supported by a CIHR CGS-D scholarship. This work was supported by the Natural Sciences and Engineering Research Council of Canada, Grants RGPIN-2020-06636 and RGPAS-2020-00014 (AE). Crystal structures solved in this work were funded in whole or in part with US federal funds from the National Institute of Allergy and Infectious Diseases, National Institutes of Health, Department of Health and Human Services, under contract No. HHSN272201700060C (Center for Structural Genomics of Infectious Diseases (CSGID); http://csgid.org). Proteomics was performed at the Network Biology Collaborative Centre at the Lunenfeld-Tanenbaum Research Institute, a facility supported by Canada Foundation for Innovation funding, by the Ontario Government, and by Genome Canada and Ontario Genomics (OGI-139), and by a Canadian Institutes for Health Research Foundation grant to ACG.

## CONFLICT OF INTEREST

The authors declare no conflict of interest.

## TABLE AND FIGURE LEGENDS

**Figure S1.**
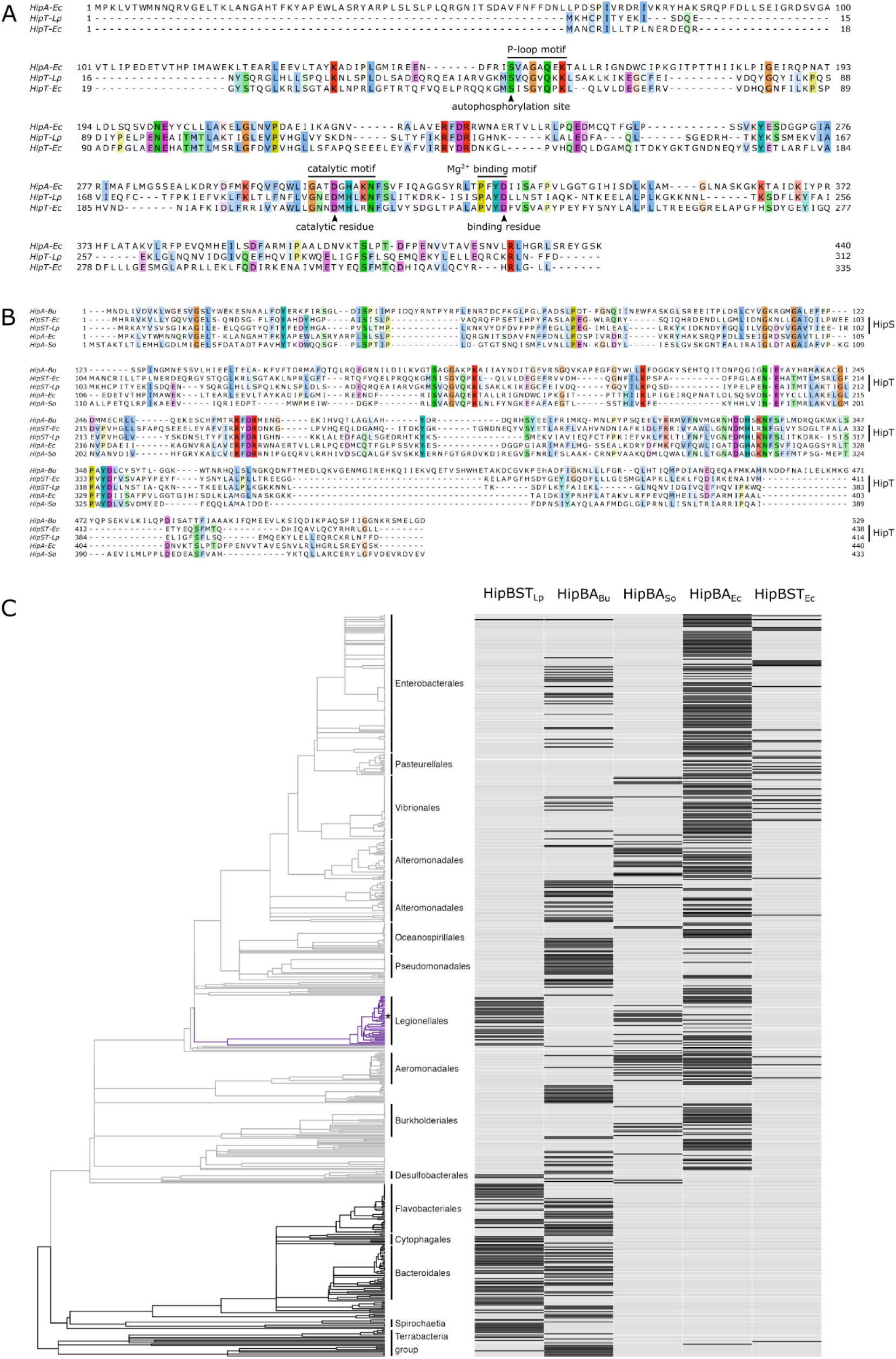
Conservation and divergence among the Hip toxins. **(A)** Alignments of HipA_Ec_ (NP_416024.1), HipT_Lp_ (AAU28431), and HipT_Ec_ (WP_001262465.1) constructed with MUSCLE. Residues are coloured with the Clustal X colour scheme implemented in Jalview (>33% identity) (76). Conserved residues and motifs are indicated. **(B)** Comparison of the seed sequences used to search for HipBA and HipBST homologs. Alignments of HipA _Bu_ (WP_149924064.1), HipA_Ec_ (NP_416024.1), HipA_So_ (AAN53784.1), HipST_Ec_ (WP_001346664.1, WP_001262465.1), and HipST_Lp_ (AAU28430, AAU28431) constructed with MUSCLE. Residues are coloured with the Clustal X colour scheme implemented in Jalview (>40% identity). **(C)** Distribution of HipBA and HipBST TA systems across diverse bacterial taxa (as in Figure 1D), but without any filtering of species based on genome completeness. The bacterial phylogeny was retrieved from TimeTree for all species containing at least one system in our homology search and the presence of each system homolog is indicated for each species (data available in Table S6). Systems are ordered by similarity of taxonomic distribution. The Pseudomonadota phylum is coloured light grey, the Legionellales order is coloured purple, and *L. pneumophila* is indicated with an asterisk.

**Figure S2.**
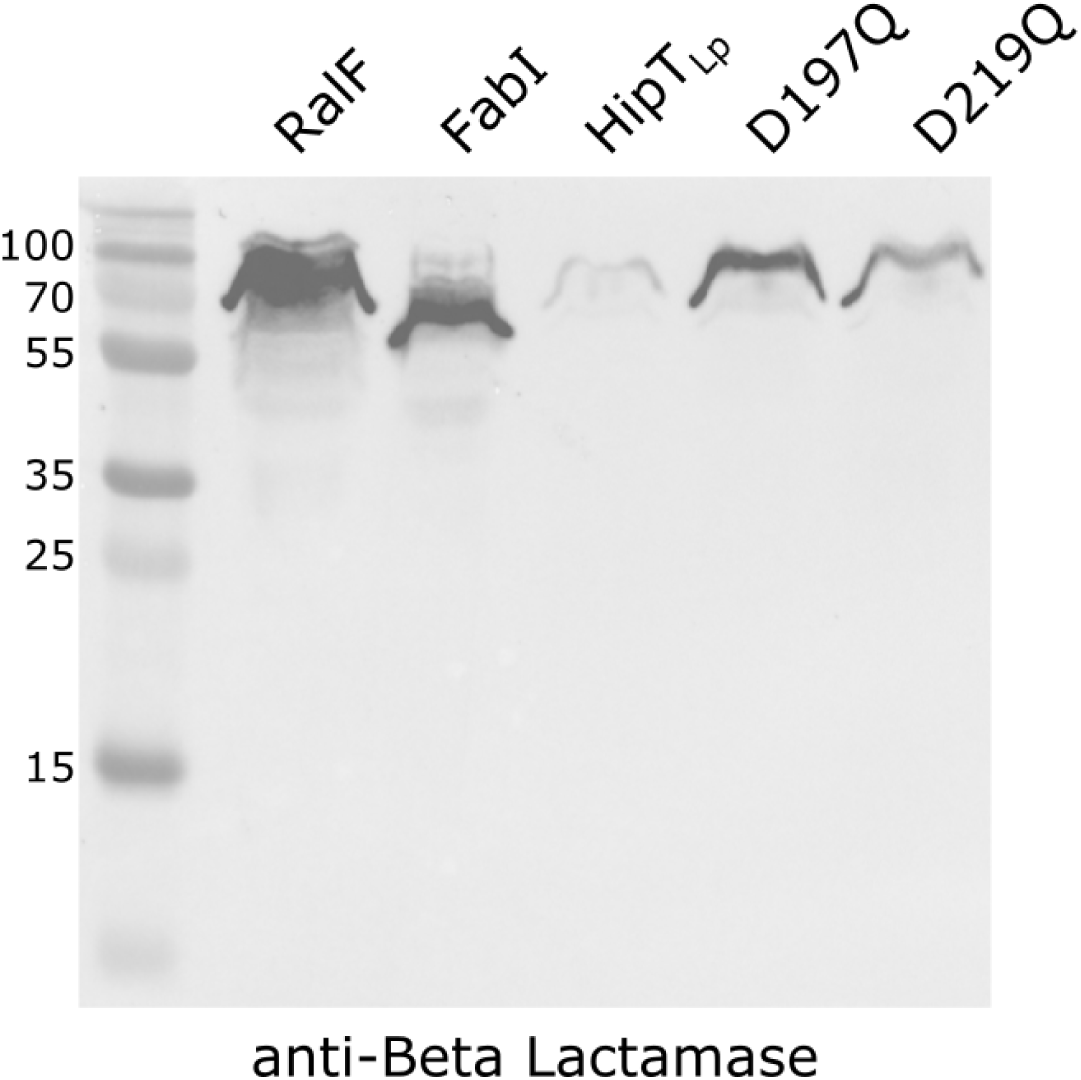
TEM-1 fusion proteins are expressed in *L. pneumophila* cells prior to translocation assays. Western blot of TEM-1 fusion protein expression in *L. pneumophila* Lp02 cells using an anti-Beta Lactamase antibody (Abcam 12251). Fusion protein expression from the pXDC61 vector was induced with IPTG (500 μM) for 3 hr. Just prior to infection of U937 monolayers, samples of induced cells were harvested for immunoblotting. The predicted molecular weight of the TEM-1 β-lactamase is approximately 31.5 kDa.

**Figure S3.**
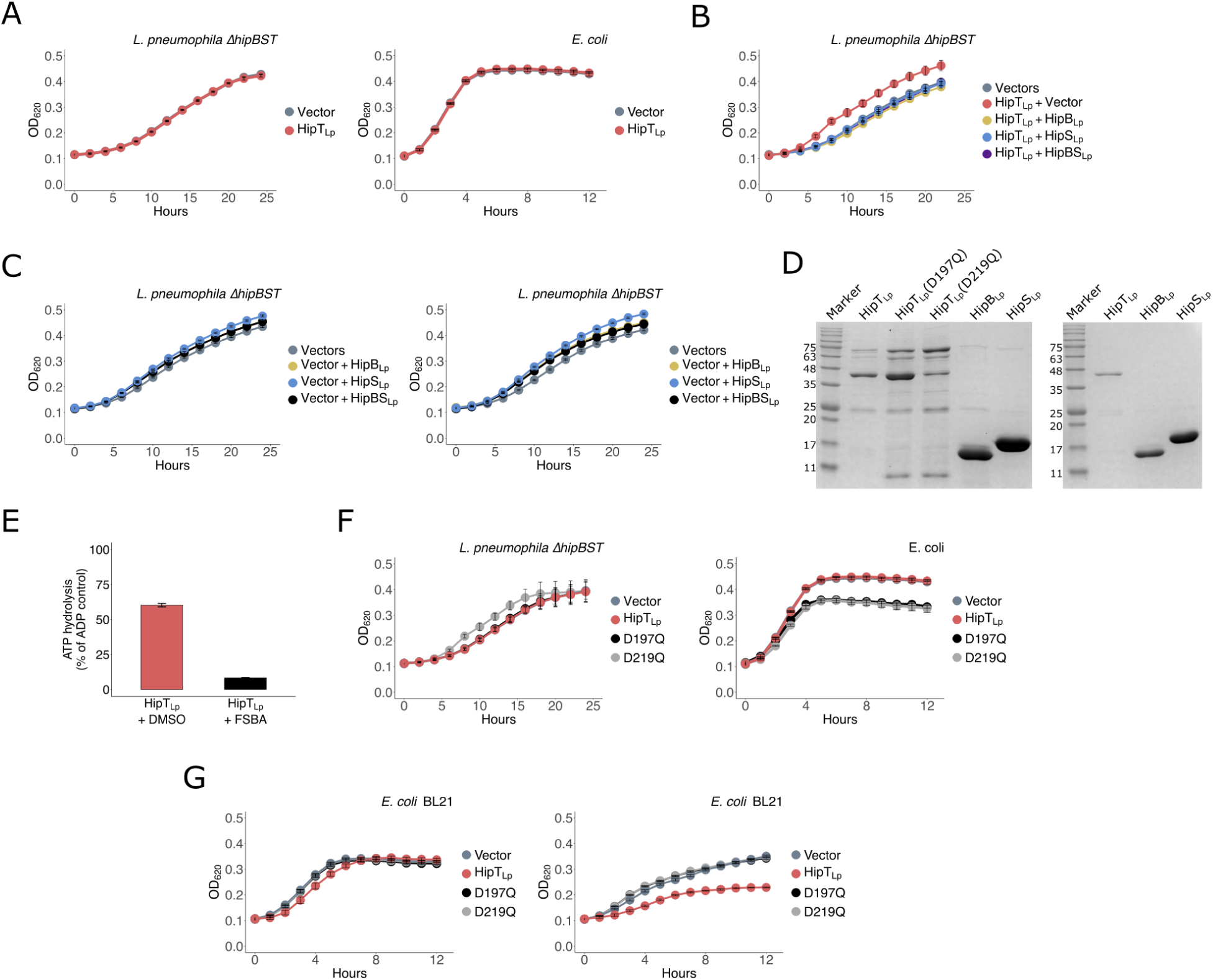
HipBST_Lp_ is a functional tripartite toxin-antitoxin system. **(A) Uninduced controls of HipT_Lp_** expression vectors in *L. pneumophila* Δ*hipBST* (pJB1806) and *E. coli* TOP10 (pBAD18) cells. Expression was repressed with 1% glucose for all constructs. **(B)** Uninduced controls of strains co-expressing HipT_Lp_ with HipS_Lp_ and HipB_Lp_ in *L. pneumophila* Δ*hipBST*. Expression was repressed with 1% glucose for all constructs. **(C)** Expression of HipB_Lp_, HipS_Lp_, and HipBS_Lp_ from pNT562 in *L. pneumophila* Δ*hipBST* cells. Uninduced controls repressed with 1% glucose (left) and cultures induced with 100 μM IPTG (right) are shown. All strains also carried the pJB1806 empty vector. **(D)** N-terminal His_6_-SBP-tagged HipB_Lp_, HipS_Lp_, and HipBST_Lp_ (wild-type and mutants) purified with nickel affinity chromatography (left gel). Proteins after secondary purification with size exclusion chromatography (right gel). Purified proteins were analyzed by SDS-PAGE and stained with Coomassie dye. **(E)** ADP-Glo kinase assay performed with purified His_6_-SBP-tagged HipT_Lp_ and the kinase inhibitor FSBA or a DMSO control. The reactions were incubated at 37°C for 30 min. Data shown are the mean ± standard deviation of 2 technical replicates and are representative of 2 independent experiments. **(F)** Uninduced controls of two HipT_Lp_ catalytic mutants (D197Q, D219Q) in both *L. pneumophila* Δ*hipBST* and *E. coli* (TOP10) cells. Expression was repressed with 1% glucose for all constructs. **(G)** Expression of wild-type HipT_Lp_ and two catalytic mutants (D197Q, D219Q) from the pJB1806 vector in BL21 cells. Uninduced controls repressed with 1% glucose (left) and cultures induced with 100 μM IPTG (right) are shown. All growth curves show the mean ± the standard deviation of 3 biological replicates. Data are representative of 3 independent experiments.

**Figure S4.**
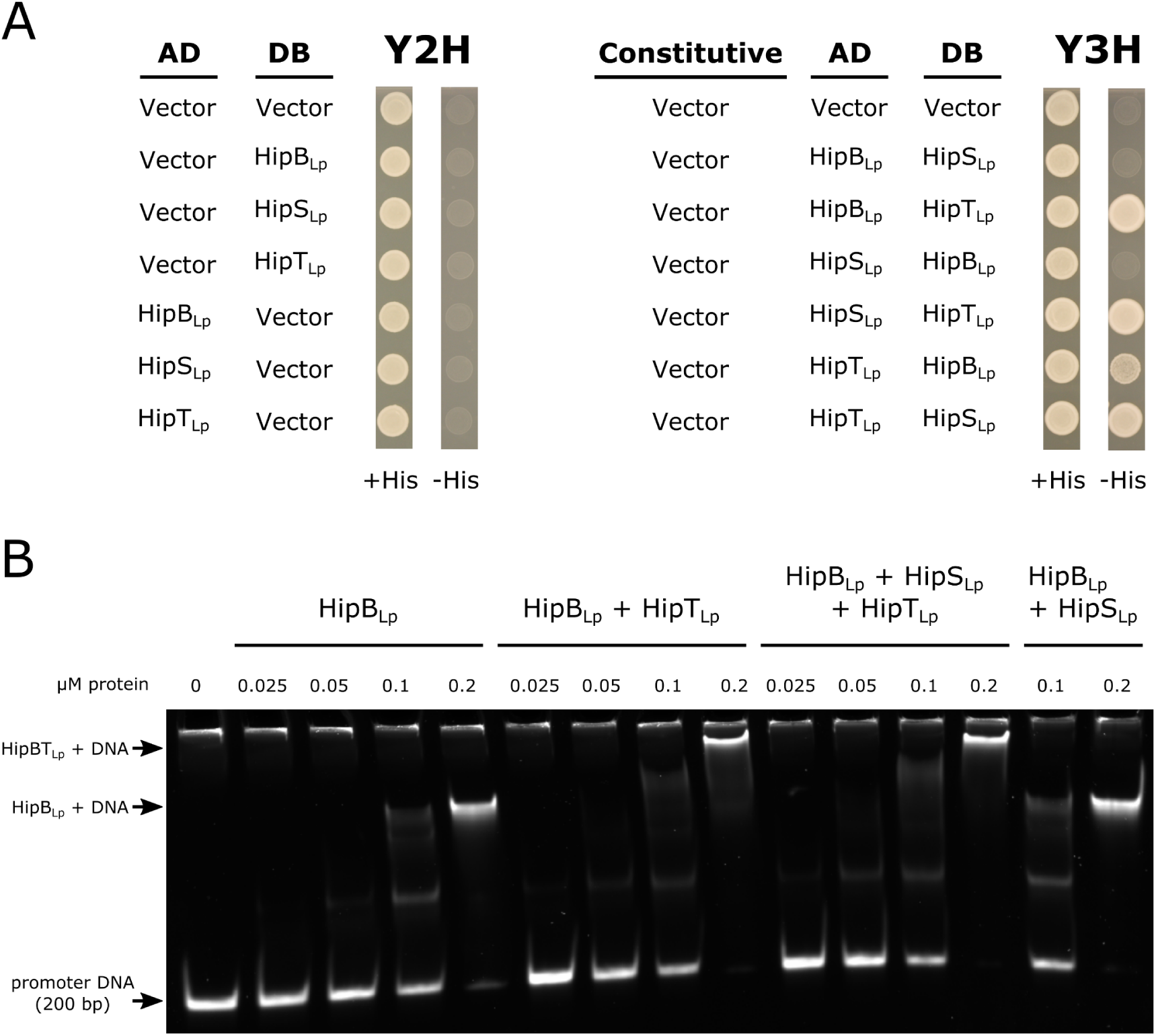
The HipBST_Lp_ system has the capacity for complex binary and ternary regulatory dynamics. **(A)** Empty vector controls for yeast two-hybrid (Y2H) experiments testing for binary physical interactions in the HipBST_Lp_ tripartite proteins. Genes cloned into the pDEST-AD and pDEST-DB Y2H vectors are indicated, and representative images are shown of *S. cerevisia*e Y8800 growth in the presence (+His) and absence (-His) of histidine. Yeast two-hybrid experiments were also performed with a third protein constitutively expressed from the pAG416 vector (Y3H). The empty pAG416 vector was used for these experiments as a control. **(B)** Electrophoretic mobility shift assay performed with recombinant purified His_6_-SBP-tagged HipBST_Lp_ proteins at equimolar increasing concentrations (indicated) and a 200 bp DNA fragment encompassing the region upstream (promoter) of the *hipBST_Lp_* locus. The gel was stained with SYBR Green and protein-DNA complexes are indicated.

**Figure S5.**
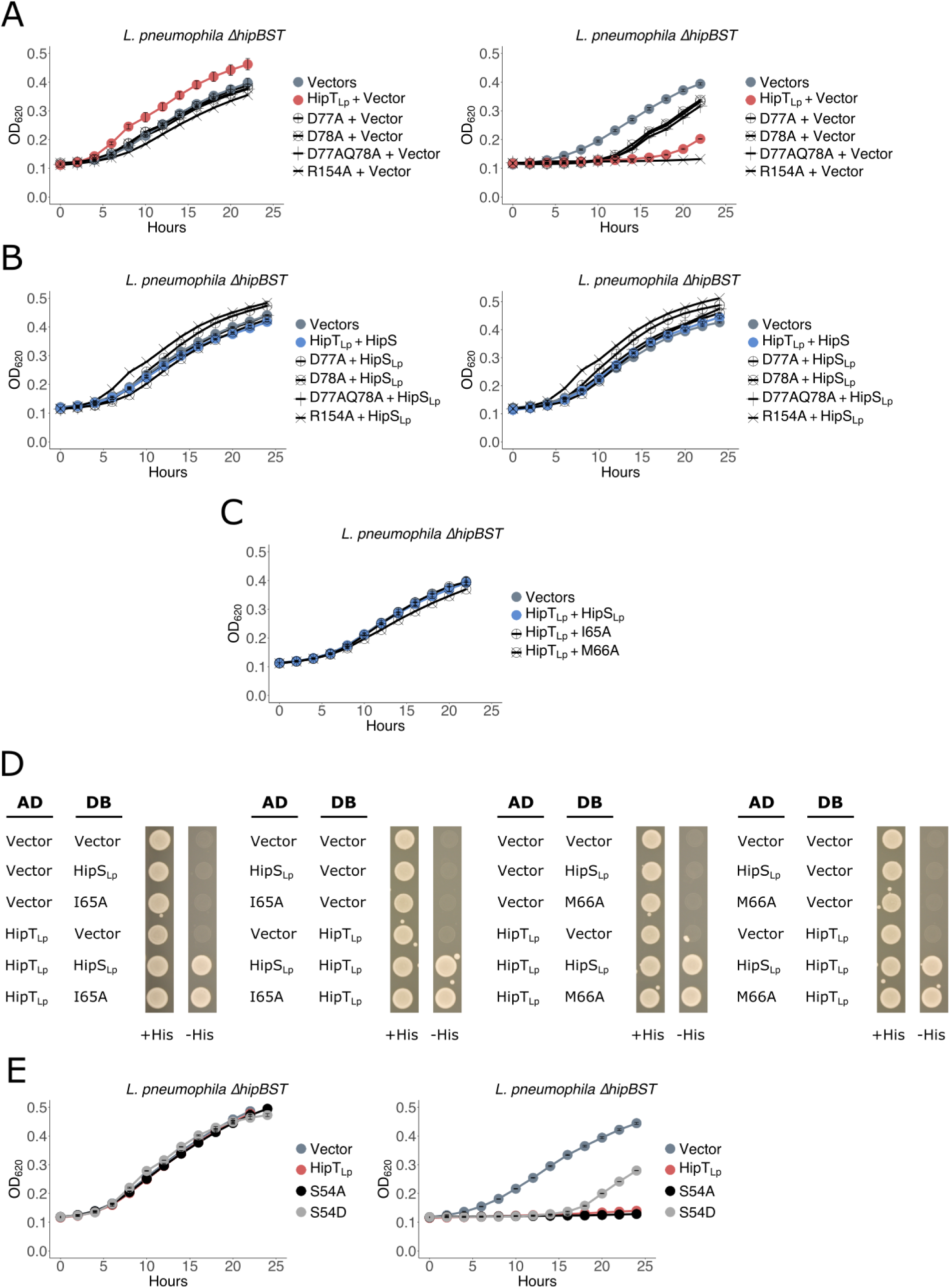
Analysis of key residues involved in HipT_Lp_ neutralization by HipS_Lp_ and autophosphorylation. **(A)** Growth assays of *L. pneumophila* Δ*hipBST* expressing HipT_Lp_ (pJB1806) with mutations in residues predicted to be important to the HipS_Lp_-HipT_Lp_ interaction interface. Experiments were performed to test whether these mutations affected HipT_Lp_ activity. Uninduced controls (left) were repressed with 1% glucose and cultures were induced with 100 μM IPTG (right). All strains also carried the pNT562 empty vector. **(B)** Growth assays of *L. pneumophila* Δ*hipBST* expressing HipT_Lp_ (pJB1806) with mutations in residues predicted to be important to the HipS_Lp_-HipT_Lp_ interaction interface. Mutants were co-expressed with HipS_Lp_ (pNT562). Uninduced controls (left) were repressed with 1% glucose and cultures were induced with 100 μM IPTG (right). **(C)** Uninduced controls (repressed with 1% glucose) for *L. pneumophila* Δ*hipBST* co-expressing HipT_Lp_ (pJB1806) with HipS_Lp_ (pNT562) bearing mutations to two residues (I65A, M66A) predicted to be important to the HipS_Lp_-HipT_Lp_ interaction interface. **(D)** Yeast two-hybrid experiments testing for binary physical interactions between HipT_Lp_ and HipS_Lp_. Both wild-type HipS_Lp_ and HipS_Lp_ bearing mutations to two residues (I65A, M66A) predicted to be important to the HipS_Lp_-HipT_Lp_ interaction interface were tested. Genes cloned into the pDEST-AD and pDEST-DB Y2H vectors are indicated, and representative images are shown of *S. cerevisia*e Y8800 growth in the presence (+His) and absence (-His) of histidine. Experiments were performed in duplicate. **(D)** Growth assays of *L. pneumophila* Δ*hipBST* expressing HipT_Lp_ (pJB1806) with phosphomimetic (S54D) and phosphoablative (S54A) mutations in its P-loop serine (S54). Uninduced controls (left) were repressed with 1% glucose and cultures were induced with 100 μM IPTG (right). All growth curves show the mean ± the standard deviation of 3 biological replicates. Data are representative of 3 independent experiments.

**Figure S6.**
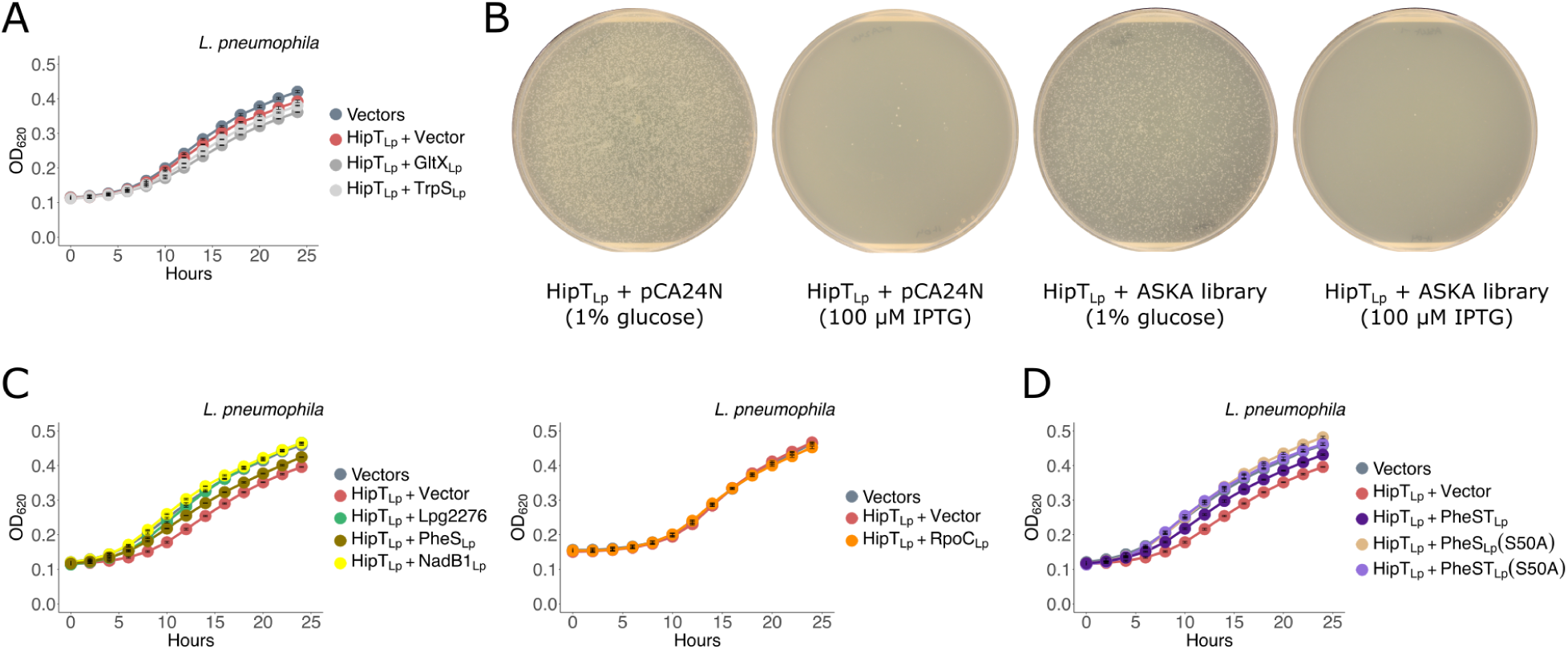
HipT_Lp_ does not target a previously characterized substrate of HipT or HipA toxins. **(A)** Uninduced controls of growth assays of *L. pneumophila* co-expressing HipT_Lp_ (pJB1806) with GltX_Lp_ or TrpS_Lp_ (pNT562). Protein expression was repressed with 1% glucose. **(B)** Example image of transformation plates from one replicate of the HipT_Lp_ rescue screen with the *E. coli* ASKA genomic library. The ASKA library was pooled and electroporated into BL21-GOLD (DE3) cells containing *hipT_Lp_* cloned into the pCDF1-b expression vector. Transformants were plated on solid media containing either IPTG (100 μM) for gene expression or 1% glucose for repression. As a control, the empty vector pCA24N was transformed in an equivalent manner to the pooled library. Library transformations were performed a minimum of three times for each experiment. Individual screening experiments were repeated 3 times. **(C)** Uninduced controls (repressed with 1% glucose) for *L. pneumophila* cells co-expressing HipT_Lp_ (pJB1806) with phosphoproteomic candidates (pNT562) that were either essential, or highly conserved in both *Legionella* and conserved in *E. coli*. **(D)** Uninduced controls (repressed with 1% glucose) for co-expression of HipT_Lp_ with PheST_Lp_ or phosphomimetic mutants of PheS_Lp_(S50A) in *L. pneumophila* cells. Growth curves show the mean ± the standard deviation of 3 biological replicates. Data are representative of 3 independent experiments.

**Figure S7.**
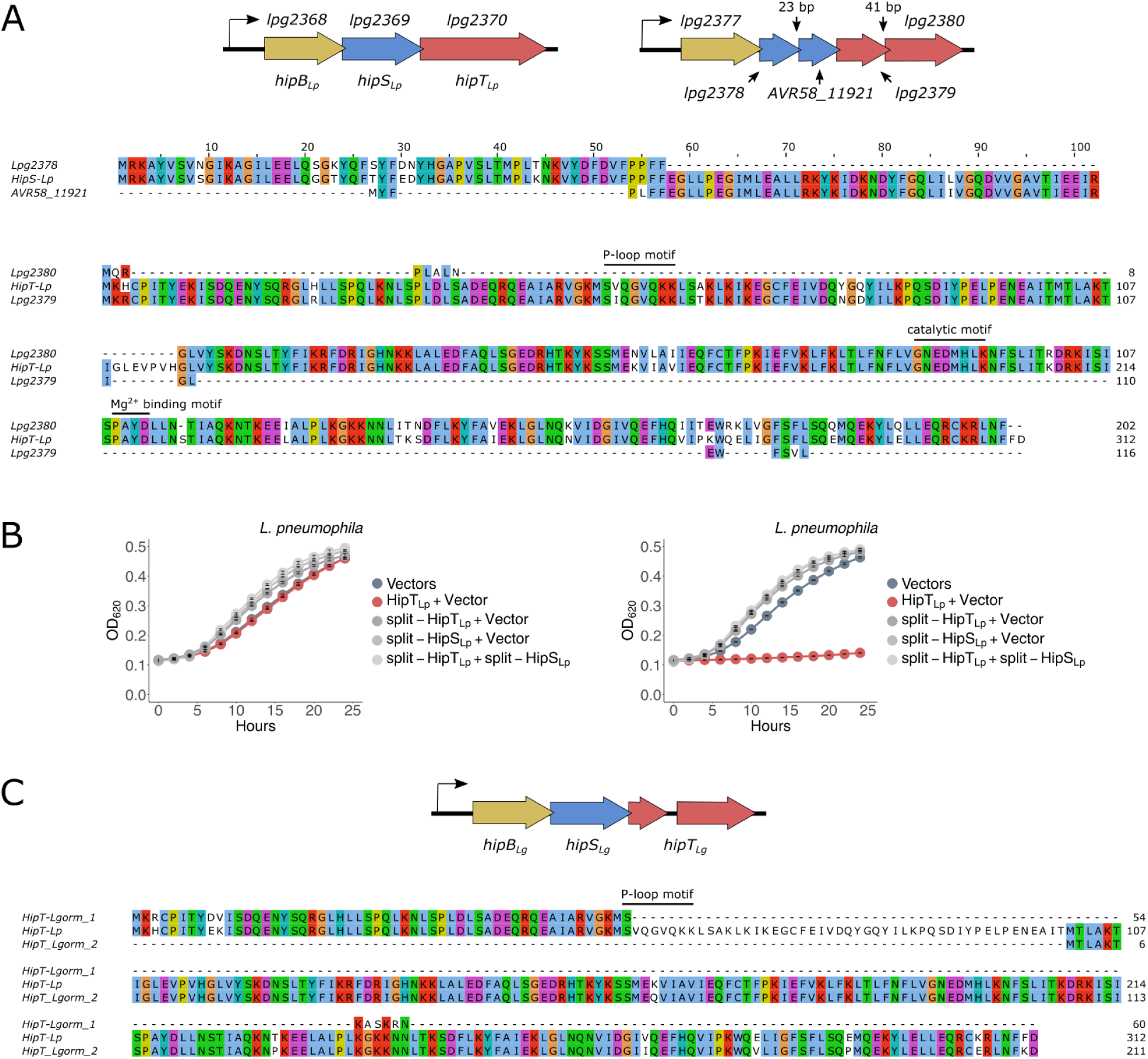
Divergent architectures suggest evolution and decay in *Legionella* HipBST systems. **(A)** Top: schematic comparing *hipBST_Lp_* with a homologous pentapartite *hipBST* locus in the *L. pneumophila* genome downstream of the *hipBST_Lp_* locus. The pentapartite module contains a split-*hipS_Lp_* and split-*hipT_Lp_* architecture (labelled). Middle: MUSCLE alignment of the split-HipS protein sequences (Lpg2378, AVR58_11921) with HipS_Lp_. Conserved residues are coloured with the Clustal X colour scheme implemented in Jalview. Bottom: MUSCLE alignment of the split-HipT proteins (Lpg2379, Lpg2380) with HipT_Lp_. Conserved motifs are labelled. Conserved residues are coloured with the Clustal X colour scheme implemented in Jalview (> 40%). **(B)** Growth assays of *L. pneumophila* cells expressing split-HipT_Lp_ (pJB1806) and split-HipS_Lp_ (pNT562), either alone or in combination. Uninduced controls (left) were repressed with 1% glucose and cultures were induced with 100 μM IPTG (right). All growth curves show the mean ± the standard deviation of 3 biological replicates. Data are representative of 2 independent experiments. **(C)** Top: schematic of a homologous quadripartite *hipBST* locus in *Legionella gormanii*, which encodes a split-*hipT* fragment that is truncated at the P-loop motif due to a frameshift. A MUSCLE alignment of both split-HipT fragments in *L. gormanii* with HipT_Lp_ is shown. Conserved residues are coloured with the Clustal X colour scheme implemented in Jalview.

**Figure S8.**
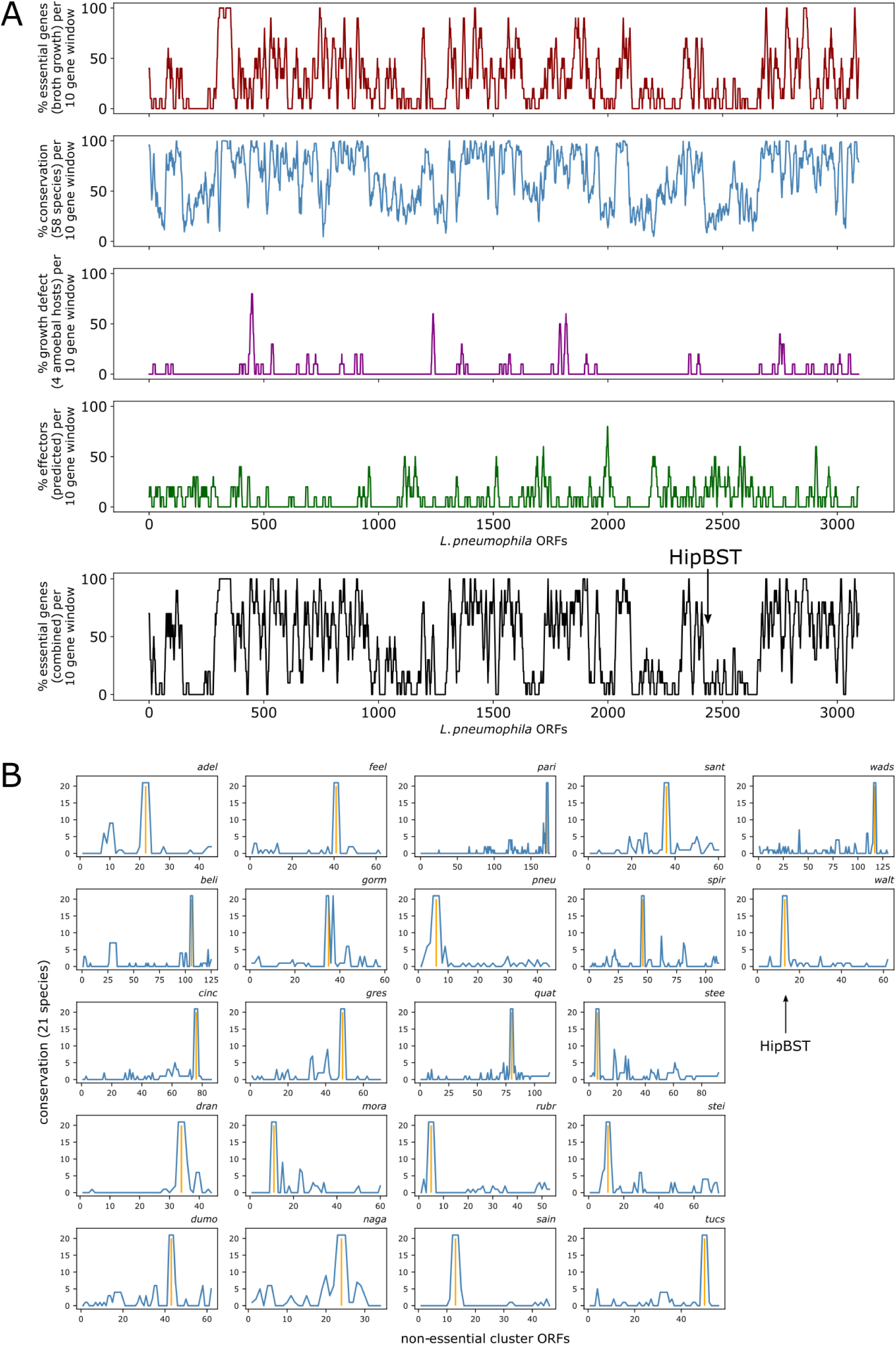
HipBST_Lp_ is associated with accessory genomic clusters of predicted non-essential genes across *Legionella* species. **(A)** The upper four panels display essentiality (broth growth) (17, 18), conservation (across 58 *Legionella* species), host replication defects (4 amoebal species) (18), and effector predictions (39) for every ORF in the *L. pneumophila* genome. Data are displayed as 10 gene sliding windows. The bottom panel displays the combined predictions for ORF conservation and essentiality (broth growth and host replication defects) across the *L. pneumophila* genome as 10 gene sliding windows. Troughs indicate accessory genomic regions of non-essential genes. The location of HipBST_Lp_ is indicated. **(B)** Accessory genomic clusters of predicted non-essential genes containing HipBST_Lp_ homologs are shown for 22 *Legionella* species. Four letter abbreviated species names are indicated for each cluster. For every ORF in each cluster (x-axis), conservation in the other 21 *Legionella* species clusters is shown (y-axis). The location of the HipBST homolog in each cluster is indicated with a vertical orange line and labelled for one cluster as an example.

## Notes

https://massive.ucsd.edu/ProteoSAFe/static/massive.jsp

### Competing Interest Statement

The authors have declared no competing interest.

